# The impact of PrsA over-expression on the *Bacillus subtilis* transcriptome during fed-batch fermentation of alpha-amylase production

**DOI:** 10.1101/2022.04.05.487122

**Authors:** Adrian S Geissler, Line D Poulsen, Nadezhda T Doncheva, Christian Anthon, Stefan E Seemann, Enrique González-Tortuero, Anne Breüner, Lars J Jensen, Carsten Hjort, Jeppe Vinther, Jan Gorodkin

## Abstract

The production of the alpha-amylase (AMY) enzyme in *Bacillus subtilis* at a high rate leads to the accumulation of unfolded AMY, which causes secretion stress. The over-expression of the PrsA chaperone aids the enzyme folding and reduces stress. To identify affected pathways and potential mechanisms involved in the reduced growth, we analyzed the transcriptomic differences during fed-batch fermentation between a PrsA over-expressing strain and a control in a time-series RNA-seq experiment. We observe transcription in 542 previously un-annotated regions, of which 234 had significant changes in expression levels between the samples. Moreover, 1,791 protein-coding sequences, 80 non-coding genes, and 20 riboswitches overlapping UTR regions of coding genes had significant changes in expression. Via gene-set over-representation analysis of the differentially expressed genes, we identified putatively regulated biological processes; overall the analysis suggests that the PrsA over-expression affects ATP biosynthesis activity, amino acid metabolism, and cell wall stability. The investigation of the protein interaction network points to a potential impact on cell motility signaling. We discuss the impact of these highlighted mechanisms for reducing secretion stress or detrimental aspects of PrsA over-expression during AMY production.

## 1 Introduction

*Bacillus subtilis* is a powerhouse for enzyme production in biotech industries (Schallmey et al., 2004; van Dijl and Hecker, 2013; Hohmann et al., 2016a). Amylases are a specific class of enzymes that *B. subtilis* can produce commercially (Schallmey et al., 2004). The amylase enzyme, in particular the alpha-amylase (AMY), is a digestive enzyme (EC 3.2.1.1) that degrades starch molecules. Therefore, AMY is often an active component in laundry detergent for removing sticky stains from cloths. For a successful AMY production and subsequent recovery, a host organism needs to both express and secrete AMY proteins in a biologically active form at a high rate (Spinnler, 2021). However, a major issue for commercial production is that the protein folding system of the cell is overwhelmed by the high rate of synthesis, unless the strains used for production are genetically modified (Kontinen and Sarvas, 1993).The accumulation of unfolded AMY proteins causes stress that requires a bacterial cell to physiologically adapt to survive (Storz and Hengge, 2010). The Sec secretion pathway secrets AMY co-translationally (Ling Lin Fu et al., 2007). Therefore, unfolded AMY is extracellular, such that the corresponding stress signal triggers the heat shock response (Westers et al., 2004, 2006; Storz and Hengge, 2010; Lim and Gross, 2014; Yan and Wu, 2019). The simplified mechanism of this stress response has two components as follows (Westers et al., 2004, 2006; Storz and Hengge, 2010; Lim and Gross, 2014; Yan and Wu, 2019): First, the membrane-bound CssS receptor transduces the stress signal by phosphorylating CssR. Second, the phosphorylated CssR activates transcription of the two proteases HrtA and HrtB, which degrade unfolded proteins and alleviate the stress condition. Further, stress responses are intertwined with additional regulation in the core energy metabolism (Storz and Hengge, 2010), and such stress responses upregulate flagellar cell motility in order for a cell to physically escape the stress causing location (Helmann et al., 1988; Marquez et al., 1990; Yan and Wu, 2019). For instance, the level of cell motility is boosted by a low level of phosphorylated DegU (Kobayashi, 2007; Verhamme et al., 2007; Gupta and Rao, 2014), which is part of the core stress regulating DegU-DegS two-component system (Storz and Hengge, 2010; Laub, 2014). Nevertheless, these stress alleviating mechanisms can be opposed to the objective of achieving a high AMY yield: (i) The proteolytic degradation of AMY reduces yields and (ii) a low phosphorylation level of DegU down-regulates AMY expression (Gupta and Rao, 2014).

A state-of-the-art approach, which prevents the yield detrimental impact of the secretion stress response, is the over-expression of PrsA (Vitikainen et al., 2001; Quesada-Ganuza et al., 2019). Although the over-expression of PrsA reduces secretion stress by aiding AMY folding, it also has detrimental impacts such as hampered cell growth and even cell lysis (Vitikainen et al., 2001; Quesada-Ganuza et al., 2019). These detrimental phenotypes might be caused by protein-protein interactions of specific PrsA protein domains with still unknown partner proteins (Quesada-Ganuza et al., 2019). Another unknown aspect of PrsA over-expression is its impact on the bacterial transcriptome, particularly during industrial fed-batch fermentation. The adaptation to glucose metabolism from maltose metabolism has a global impact on half of all transcriptional regulators even though both carbons are preferred by *B. subtilis* (Buescher et al., 2012). Thus, we would assume a substantially larger global impact on the transcriptome for the extreme secretion stress during PrsA over-expression (Quesada-Ganuza et al., 2019). We consider our assumption to be further supported by the large number of over a hundred proteins that require regulation to adapt bacterial motility (see above concerning stress) (Rajagopala et al., 2007). Furthermore, a pure protein-coding gene focus ignores the essential role regulatory small RNA (sRNA), RNA chaperones, and non-coding RNA (ncRNA) have in facilitating physiological changes impacting the entire cell during stress responses (Storz and Hengge, 2010). General stress regulatory mechanisms have been investigated in public datasets (Arrieta-Ortiz et al., 2015); however, metabolic and stress pathways undergo complex temporal adaptations (Hahne et al., 2010; Otto et al., 2010). Thus, both temporally resolved and condition-specific gene expression levels are needed to study stress pathways. Specifically for secretion stress during *B. subtilis* AMY fed-batch fermentation, no such dataset exists to our knowledge.

Here, we conducted fed-batch fermentation of two commercial *B. subtilis* strains. Both strains produce an AMY and are isogenic, except that one of them over-expresses PrsA. We studied the transcriptome during fermentation at 6 timepoints with RNA-seq and analyzed the expression levels of both known coding and non-coding annotations, but also of potential novel transcribed, yet un-annotated regions. We complemented the differential expression analysis with a network analysis of known protein-protein interactions (PPI). This study found significant changes in gene expression levels between the studied strains for genes in the ATP biosynthesis and cell motility biological processes. Further, the network analysis hints at mechanisms relating to competence transformation and cell motility that might be candidates for further tuning of AMY secretion yields.

## 2 Materials and Methods

### 2.1 Strains and fed-batch fermentation

The overall experimental setup is as previously described in (Geissler et al., 2022). In summary: *B. subtilis* strain 168 *ΔspoIIAC ΔamyE Δapr ΔnprE ΔsrfAC* was maintained at 4 °C on LBGG medium. The AMY JE1 (sequence label *je1zyn* in (Geissler et al., 2022)) was inserted by Splicing by Overlapping Extension (SOE) linear recombinant transformation, together with the commercial *sigA* promoter sequence P4199 and chloramphenicol marker, in the *pel* locus. The PrsA over-expressing strain (referred to as ‘+prsA’ strain) had the insert by SOE of P4199, *prsA*, and spectinomycin marker in the *amyE* locus. A control strain did not have the *prsA* insert. After inoculation on SSB4 agar at 37 °C, transfer on M-9 medium, sucrose 2M fed-batch fermentations were conducted in proprietary 5L tanks at 38 C. Fermentations were run in triplicates for 5 days. The selected replicate size allows detecting significant logFC in expression of at least ±0.5 magnitude, as determined in benchmarks (Schurch et al., 2016). Samples were taken at 6 timepoints: 21 h, 26 h, 45 h, 71 h, 94 h, and 118 h after fermentation started. The samples were measured in cell density (OD650), and AMY activities were measured with an in-house assay. The assay (after 1/6000 dilution) states the enzyme amount that breaks down 5.26 g starch per hour. This activity measure is proportional to the enzyme yield.

### 2.2 RNA-seq dataset

All samples were immediately mixed with 5 ml 100% ethanol and stored on dry ice. The RNA extraction and purification method is the identical phenol-chloroform protocol of (Geissler et al., 2022). RNA libraries and sequencing were conducted by BGI Hong Kong with DNBseq in single-ends of 50 bp length. RNA libraries were prepared with 3’ adapter sequence AAGTCGGAGGCCAAGCGGTCTTAGGAAGACAA and the 5’ adapter AAGTCGGATCGTAGCCATGTCGTTCTGTGAGCCAAGGAGTTG. The 36 samples (triplicates, 2 strains, 6 timepoints) were sequenced in 3 batches with technical replicates for QC (Supplementary Table S1). The computational analyses were conducted in an adapted workflow of (Geissler et al., 2022) (doi: 10.5281/zenodo.4534403), which provides a pipeline in a Snakemake framework nested in computational reproducible Anaconda environments (Koster and Rahmann, 2012). In concordance with the read quality assessment of FastQC (version 0.11.8) (Simon Andrew), any adapter contaminations were removed with Trimmomatic (version 0.39) for up to 2 seed mismatches at a minimal 10 bp sequence overlap and 30 bp palindromic overlap (Bolger et al., 2014). In a sliding window of 4 bp, reads were clipped for average PHRED score quality below 20. From the 3’ of reads, positions with quality below 3 were removed. Finally, a minimal length of 40 bp was required for filtered and cleaned reads. Reads were mapped against the respective +prsA and control genome sequence with Segemehl (version 0.3.4, default settings) (Hoffmann et al., 2009). The mapping and QC filtering statistics are in Supplementary Table S2. Expression levels of coding and non-coding annotations (see below) in the respective strains were quantified for uniquely mapping reads with featureCounts (Subread version 1.6.4, ≥50% overlaps). Annotation coordinates in the respective strains were determined by liftOver (version 377) from the reference assembly (NC_000964.3) based on a pairwise alignment with LASTZ (version 1.0.4) (Harris, 2007; Liao et al., 2014; Haeussler et al., 2019).

### 2.3 Novel potentially transcribed regions

Reference annotations of coding, non-coding RNA (ncRNA), transcripts, untranslated regions (UTRs), and RNA structures were used from the BSGatlas (version 1.1). The BSGatlas uses separate annotation entries to specify which regions of an mRNA transcript are the coding, untranslated, or potential cis-regulatory RNA structure parts. Such a distinguishment to the UTR element is advantages since cis-regulatory RNA structures can overlap coding regions. Additional 141 putative ncRNA annotations from a tiling-array study were used (which are not part of the BSGatlas) (Nicolas et al., 2012; Geissler et al., 2021). Relative to these reference annotations and all transcript and untranslated regions (UTRs) annotated in the BSGatlas, we checked our RNA-seq data for transcription signals in 1,645 un-annotated regions. The additional tiling-array annotations and un-annotated regions were determined with the R library plyranges (version 1.6.0) and GenomicRanges (version 1.38.0) combined with an overlap helper script from BSGatlas’ analysis code (doi: 10.5281/zenodo.4305872) in R (version 3.6.3) (R Development Core Team, 2008; Lawrence et al., 2013; Lee et al., 2018). Un-annotated regions shorter than 100 bp (the minimum length for >99% of the transcripts in the BSGatlas) were excluded from any further expression analysis. The expression counts for all coding/non-coding sequences and cis-regulatory RNA structures were normalized with DESeq2’s size-factor estimation (version 1.26.0) (Love et al., 2014). With respect to the downstream analysis of expression signals, we excluded the UTR annotations for improved interpretability, although we still retained all structured RNA cis-regulatory annotations. With the possible overlap between cis-regulatory RNAs and coding sequences, reads mapping within such overlaps can be counted twice during the quantification of expression. For a total of 542 un-annotated regions, we observe expression signals of normalized read counts relative to gap length of at least 4 / 50 bp (corresponds to 4 times average coverage) (Supplementary Fig S1). We chose not to narrow down the transcribed regions, because we found that a read coverage-based approach (as suggested in the workflow used in the RNA-seq dataset, last section) resulted in fragmented results (see example in Fig S9). These regions were assumed to be *novel potentially transcribed regions* (NPTRs) (see Supplementary Table S3); all other un-annotated regions were excluded from the subsequent expression analysis.

### 2.4 Differential expression analysis

The expression levels of the coding/non-coding sequences, NPTRs, and cis-regulatory RNA structures were assessed for biological reproducibility in expression counts with scatter plots (Supplementary Fig S2). The scatter plots did not indicate visually striking patterns of batch effects according to the sequencing plan (Supplementary Table S1). The principal component analysis (PCA) inspection of the top 100 most variant expressed annotations (without further diff. expression analysis) confirmed the relevance of the experimental design in the latent structure of the expression data with the principal components corresponding to the strains and time aspect (Supplementary Fig S3). Differential expression for pairwise comparisons between the strains at each of the 6 time points and within each strain along the time axis (Fig 1 C) were assessed with DESeq2’s Wald test. Similar to the analysis presented in (Geissler et al., 2022), the pairwise tests were weighted in a stage-wise procedure to guarantee an overall relative to the number of annotations: Each annotation was screened for dynamic expression with a log-ratio test against a static expression model before confirming which of the pairwise tests had changes in expression. The screening and pairwise tests included a linear factor in the regression models to account for potential batch effects. The stage-wise weighting was conducted with stageR (version 1.8.0) (Van den Berge et al., 2017) and differential expression was called for adjusted p-values < 0.01. Overall, 2,127 annotations were detected as differentially expressed (Table 1, Supplementary Table S4). Based on the z-scaled log expected mean expression levels (Supplementary Table S5), expression profiles were grouped in 10 k-means clusters (R implementation). The profiles per strain were clustered separately (one gene = two rows in the data matrix). The number of clusters was determined by the “elbow method” over the total within-cluster error curve (Supplementary Fig S4) (Thorndike, 1953).

**Figure 1.**
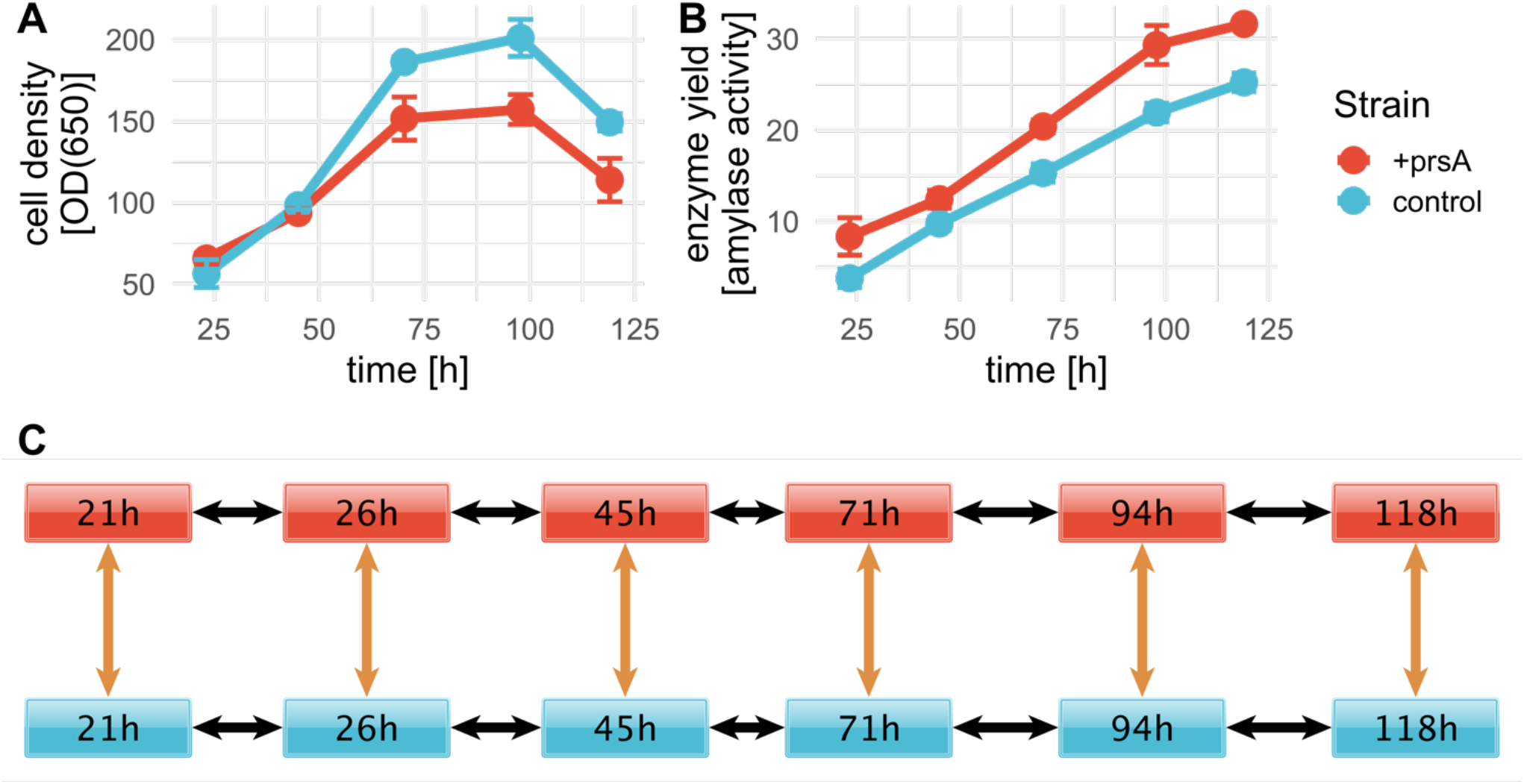
AMY fed-batch fermentations. Fed-batch fermentation was conducted in triplicates for a control strain (blue) and +prsA (red). RNA-seq samples were prepared at 6 timepoints: 21 h, 26 h, 45 h, 71 h, 94 h, and 118 h after fermentation start. Cell density and enzyme yield were measured for 5 timepoints: 23.2h, 45h, 70.2h, 97.8h, 119h.(**A)** The average cell density per strain over fermentation time was measured in optical density (OD) at 650 nm. The error bars indicate the standard deviation. **(B)** With a progressing fermentation, the yield increases. The shown yield is measured in enzyme activity (see methods “strains and fed-batch fermentation”). **(C)** For the differential expression, we investigated the significance of differential expression between the samples at 6 pairwise comparisons (orange arrows) and changes in expression over time in either strain for each pair (black arrows).

**Table 1.**
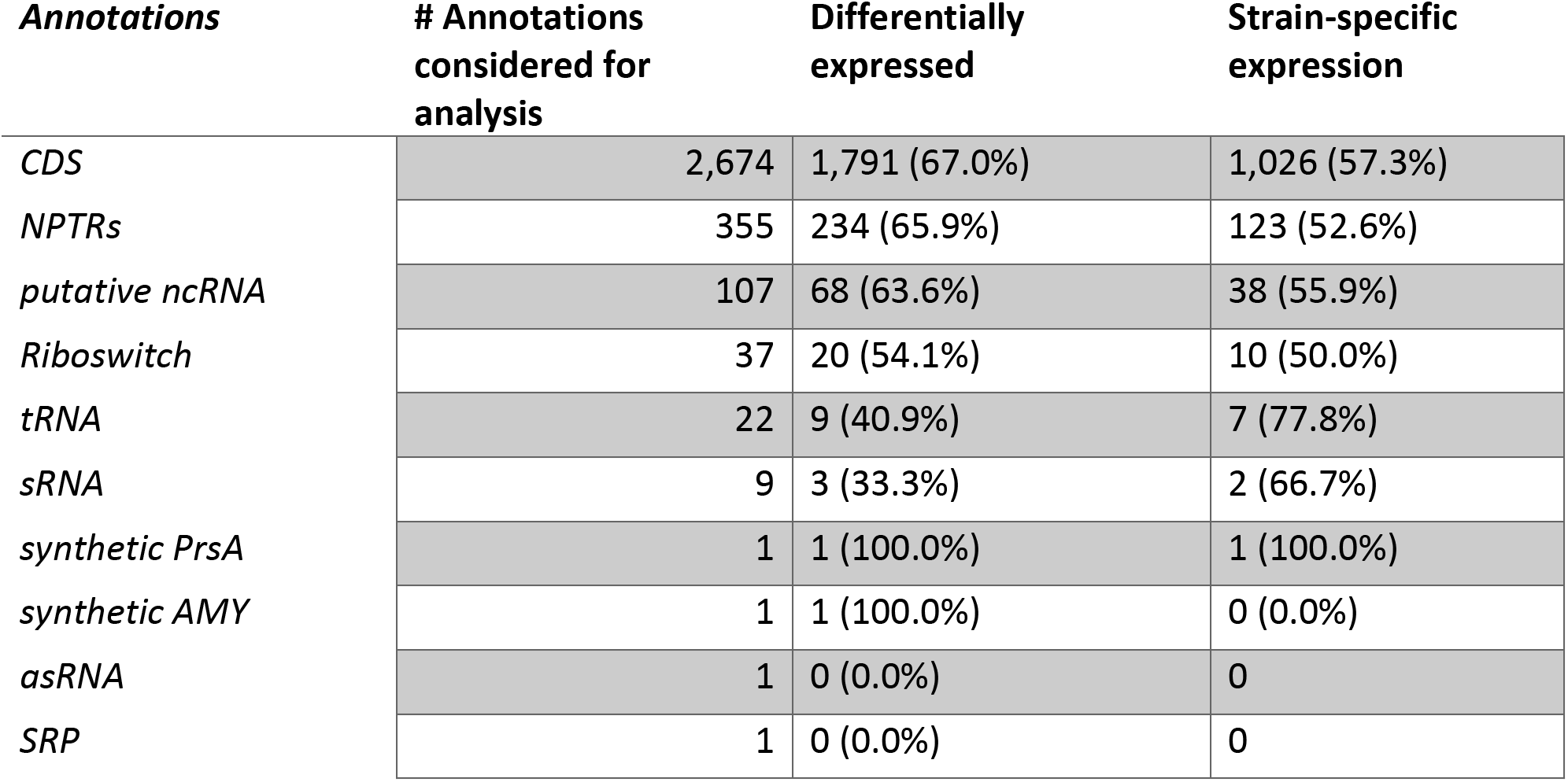
Differentially expressed annotations. For the differential expression analysis, multiple coding and non-coding annotations were considered (first column). The number of genes with minimal expression levels as determined by DESeq2’s independent filtering, which were inspected for potential differential expression, is in the second column. The number of detected differentially expressed annotations in any of the pairwise comparisons (Fig 1 C) is in the third column. The last column lists the number of annotations detected to have a significant difference in expression between the strains. The percentages provided in parenthesis are relative to the columns to the left. (Note: Only 355 of the 542 NPTRs passed the independent filtering)

### 2.5 Regulated biological processes

We investigated the set of differentially expressed genes and their up- and downregulation for over-representation in biological processes as annotated in Gene Ontology (GO) terms, which are readily available for 78.3% of coding genes (Caspi et al., 2014; Geissler et al., 2021). For each pairwise differential expression test (Fig 1 C), we inspected the set of upregulated genes (those with a positive logFC) and downregulated genes separately. The over-representation analysis was performed with topGO (version 2.37.0) (Adrian Alexa and Jörg Rahnenführer). Over-representation for the respective up- and downregulated genes was determined with a fisher test for the significance level of 0.01 relative to the background of all expressed genes, which were determined by DESeq2’s independent filtering procedure. This procedure discards the on average lowly expressed genes in order to maximize the number of differentially expressed genes (indicated by NA for p-values in Supplementary Table S4) (Love et al., 2014). The minimal term size was set to 10, and the dependencies due to GO’s hierarchy were de-correlated with topGO’s “elim” algorithm. After filtering for a minimal observed/expected ratio of magnitude 2 (between the 80 and 85^th^ percentile), p-values were adjusted for multiple testing with false discovery rate (FDR). The over-represented processes and the associated differentially expressed genes are listed in Supplementary Tables S6 and S7 and Figure S6.

### 2.6 PPI network analysis

The PPI network analysis was conducted in Cytoscape (version 3.8.2) (Shannon, 2003) for the differentially expressed protein-coding genes (both with and without significant logFC between strains). High-confidence protein associations (confidence score > 0.8) were retrieved from the STRING v11 database using stringApp (version 1.6.0) for the *B. subtilis* strain *168* (Doncheva et al., 2019; Szklarczyk et al., 2019). The resulting network was clustered with the MCL algorithm (inflation value of 2.5, confidence scores as edge weights) implemented in clusterMaker2 app (version 1.3.1) (Enright, 2002; Morris et al., 2011). The visualization of significant between strain logFCs on the network nodes was added with Omics Visualizer (version 1.3.0) (Legeay et al., 2020).

### 2.7 Global amino acid composition

In order to interpret the regulated biological processes (see above), we inspected the global amino acid compositions of all *B. subtilis* protein-coding genes. The nucleotide sequences of all coding sequences from the BSGatlas were extracted with BSgenome (version 1.54.0) (Pagès, 2021). The corresponding amino acid sequences were determined according to the bacterial genetic code with Biostrings (version 2.54.0) (Pagès et al., 2019). Here, we used only the 99.3% of the coding genes that were complete relative to their corresponding amino acid sequences; that is, they used all codons encoded in their nucleotide sequences, correctly started with methionine, and ended with a stop codon. The composition in average proportion was determined for these complete sequences (Table 3).

## 3 Results

### 3.1 Novel potentially transcribed regions

#### 3.1.1 Transcriptome analysis from RNA-seq data

To elucidate potential mechanisms of *B. subtilis* secretion stress during the production of the AMY enzyme JE1 (commercial name Natalase™) with a particular focus on PrsA over-expression, we conducted fed-batch fermentation in triplicates for two isogenic strain conditions: One control strain and one strain with PrsA over-expression (from here on referred to as +prsA). As expected from the reduced growth upon PrsA over-expression (Vitikainen et al., 2001; Quesada-Ganuza et al., 2019), the +prsA strain has a lower cell density (Fig 1 A) and higher AMY yield (Fig 1 B). To capture the transcriptome dynamics during fermentation, we took out samples for RNA-seq analysis at6 timepoints: 21 h, 26 h, 45 h, 71 h, 94 h, and 118 h after fermentation started. These timepoints correspond to two samples for the first day of fermentation and one sample per remaining day.

#### 3.1.2 Transcriptional activity for the reference annotations

In order to comprehensively investigate both the coding and non-coding RNA elements, we quantified the RNA-seq expression according to a recently developed transcript atlas for *B. subtilis* (Geissler et al., 2021). We included 141 additional annotations from a tiling-array study that was not included in the atlas due to unclear mechanism of transcription (annotations were ambiguous to whether they are independent full RNA transcript or only part thereof (Nicolas et al., 2012; Geissler et al., 2021). In the following, we refer to these annotations, together with the less well-characterized RNA elements from the atlas, as putative ncRNA. These reference annotations combine gold standard curated information, computational RNA structure biology, and transcriptomic analysis of over 100 experimental conditions (Nicolas et al., 2012; Geissler et al., 2021). Additionally, these experimental conditions suggest that still 5% of remaining un-annotated regions have evidence of expression activity (Geissler et al., 2021). Fed-batch fermentations were not part of the above-mentioned experimental conditions, such that there might be a larger potential to discover fed-batch related regions from our RNA-seq data. Consequently, we investigated our RNA-seq data for expression in such un-annotated regions.

#### 3.1.3 Novel potentially transcribed regions

There are a total of 1,645 un-annotated contiguous stretches of the genome or gaps (stranded, meaning there can be antisense located annotations) between reference annotations of length > 100bp (minimal length for 99.5% of transcripts in the atlas). We detect novel potentially transcribed regions (NPTRs) by inspecting the average RNA-seq read coverages over the entire un-annotated gap region (read counts, DESeq2 size-factor normalized, relative to the lengths). Relative to the 50 bp sequencing lengths (see methods “RNA-seq dataset”), 70% of atlas annotations were on average expressed by four reads and 30% by one read. In contrast, only 20% (542) of un-annotated regions were on average covered by four reads. This high coverage for these 542 NPTRs (Supplementary Fig S1) indicates that the NPTRs may have functional importance and that it would be relevant to include these in subsequent expression analysis (see Supplementary Table S3).

### 3.2 PrsA over-expression changes gene expression regulation of global transcriptome

#### 3.2.1 Differential expression

We assessed the impact of PrsA over-expression on the bacterial transcriptome by analyzing the expression levels of coding and non-coding sequences (see “Transcriptional activity for reference annotations” above), including the 141 additional annotations and the 542 NPTRs with DESeq2. For each region, we performed16 pairwise differential expression tests: 6 tests between the two strains on each timepoint and 2×5 tests from one timepoint to the next in both strains (Fig 1 C). Since each pairwise test corresponds to a separate hypothesis test, we used stage-wise testing to adjust for the overall false discovery rate (FDR) per annotation (Love et al., 2014; Van den Berge et al., 2017). Compared to controlling the FDR per hypothesis, the overall FDR increases statistical power and guarantees the FDR relative to the gene/annotation number, independent from the number of hypotheses (Van den Berge et al., 2017). As part of the differential expression analysis, DESeq2’s independent filtering detected about half of all coding sequences and 355 of 542 NPTRs as expressed (Love et al., 2014). At an overall FDR p-adj. ≤ 0.01, we detected differential expression for 1,793 coding sequences (67% of expressed genes), 234 NPTRs (66%), 68 putative ncRNAs (64%), 20 riboswitches (54%), 9 tRNAs (41%), and 3 sRNAs (33%) (Table 1, Supplementary Table S4). The differentially expressed coding genes include the AMY enzyme and the over-expressed PrsA. Between 50 and 78% of these biotypes had strain-specific expression patterns (significant difference for at least one of the 6 between strain tests). PrsA had strain-specific expression (as to be expected by not being inserted into the control strain’s genome). Notably, no strain-specific expression was detected for AMY.

#### 3.2.2 The regions with the highest expression changes

The strain-specific expression patterns of PrsA and the respective logFC between the two strains on all 6 timepoints were the most extreme observed in this study with logFC values up to a factor of 20 at each timepoint. Other extreme logFC values were observed for genes from operons encoding a variety of biological functions (Table 2). The NAD biosynthesis genes of the *nadABC* operon (Rodionov et al., 2008) also have extreme logFC, but they undergo both extreme up- and downregulation in the control strain with *nadA* and *nadB* being downregulated from timepoint 21h to 26h (both logFCs < −6, adj. p < 0.004) and subsequently upregulated from 26h to 45h (both logFCs ~+7, adj. p < 3e-10). Due to the secretion stress the production strains attempt to sporulate despite being unable to do so (Geissler et al., 2022). Consistently, the two sporulation genes *safA* and *coxA* were among the most extremely regulated (logFC > 6, adj. p < 2.3e-5). Other extreme logFC (< −5, adj. p < 7.31E-09) were observed for the spore killing factors *skfA* and *skfB* (González-Pastor, 2011), the sporulation controlling factor *spoIIGA* (Ramos-Silva et al., 2019), the bacitracin resistance genes *bceA* and *bceB* (Ohki et al., 2003), the for NADH during fermentation essential lactate dehydrogenase *ldh* (Cruz Ramos et al., 2000; Larsson et al., 2005), and an NPTR antisense to the gene of unknown function *ytta* (Asai et al., 2007).

**Table 2.**
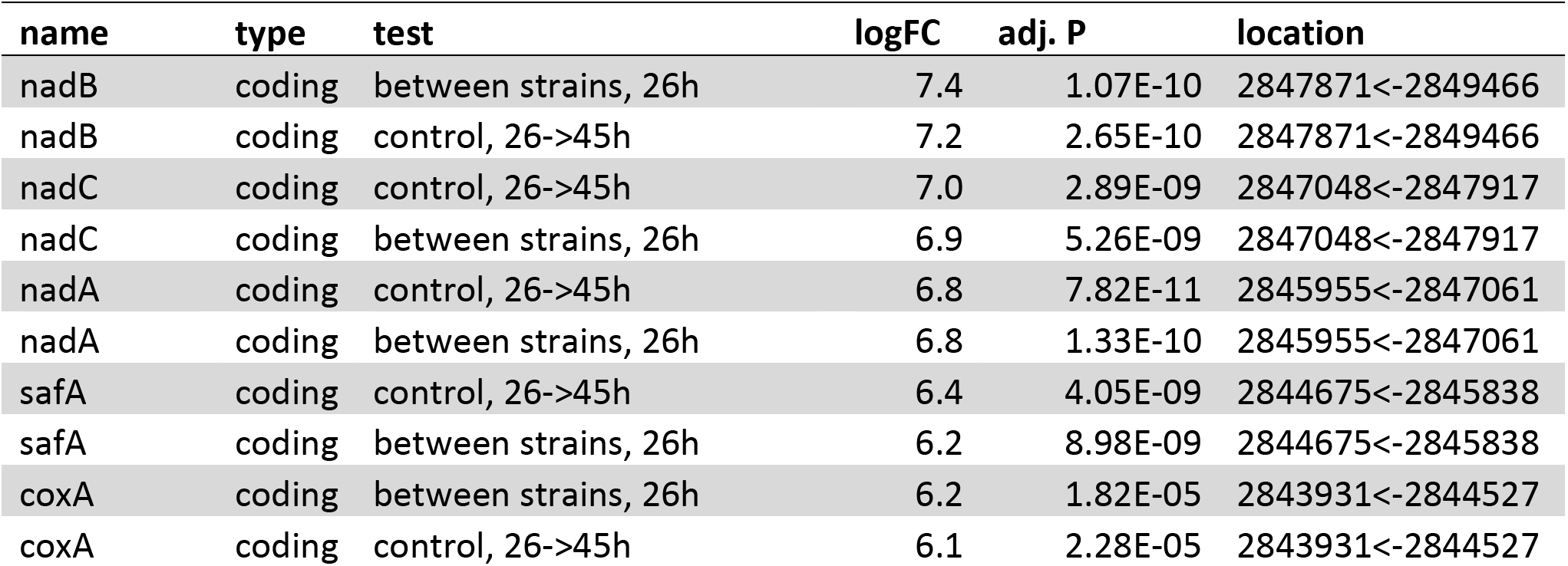
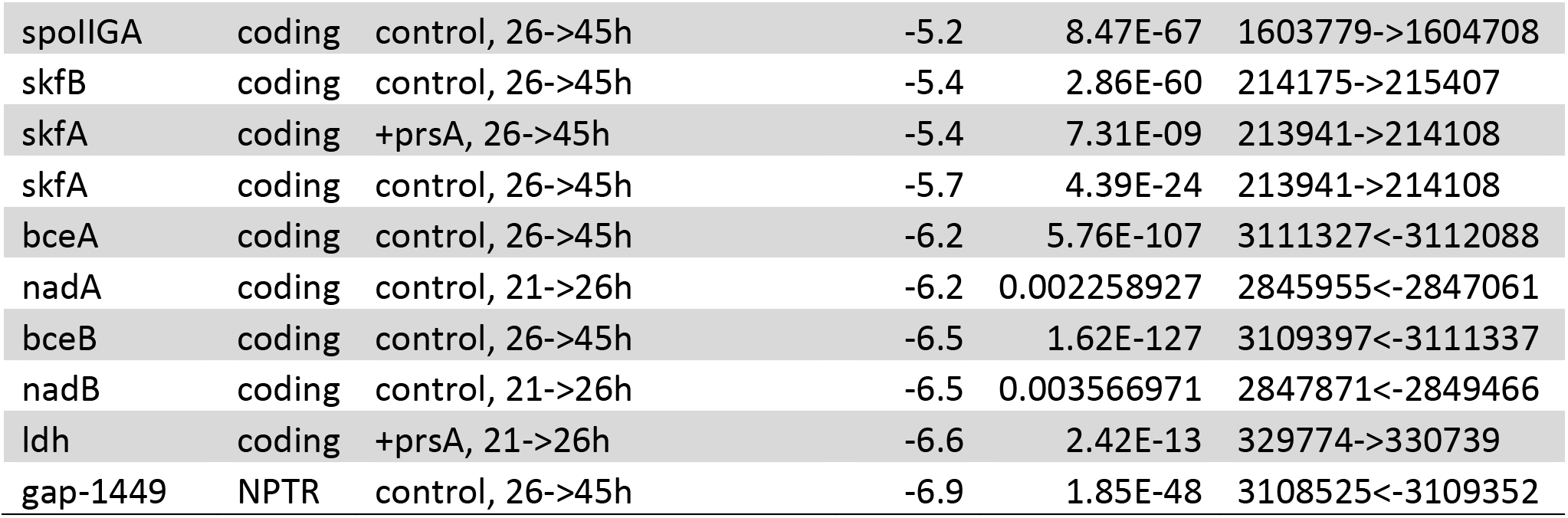
Most extreme observed logFCs. The table lists the top 10 most extreme up- and down-regulated genomic elements according to their logFC of differential expression (fourth column). prsA is excluded since it was upregulated with an approximate logFC of 20 between strains at all time points. For each genomic element, the locations (last column) are relative to the reference genome (see methods). The pairwise tests (third column) refer to the conducted differential expression analysis (Figure 1C), and the corresponding adjusted P-values are listed in the fifth column.

#### 3.2.3 Biological processes and differentially expressed genes are mutually associated

The investigation of the overall expression profiles from a k-means clustering on the average expected expression at each timepoint (Supplementary Table S5) shows marked differences in the expression dynamics between the strains (Fig 2 C). Also, all profiles indicate a substantial shift in dynamics between timepoints 45-71 h, during which the cell population increased the most (Fig 1 A): For instance, profiles 4 and 5 drop in expression levels at that timepoint but recover and even exceed the starting expression level whereas profiles 7 and 8 have drastically downregulated expression at that timepoint and do not recover (Fig 2 B). Genes and other biotypes with strain-specific expression patterns had predominately different expression profiles between the strains, whereas those without strain-specific expression had the same (Supplementary Fig S5). Therefore, *B. subtilis* regulates gene expression both timepoint- and strain-specifically.

**Figure 2.**
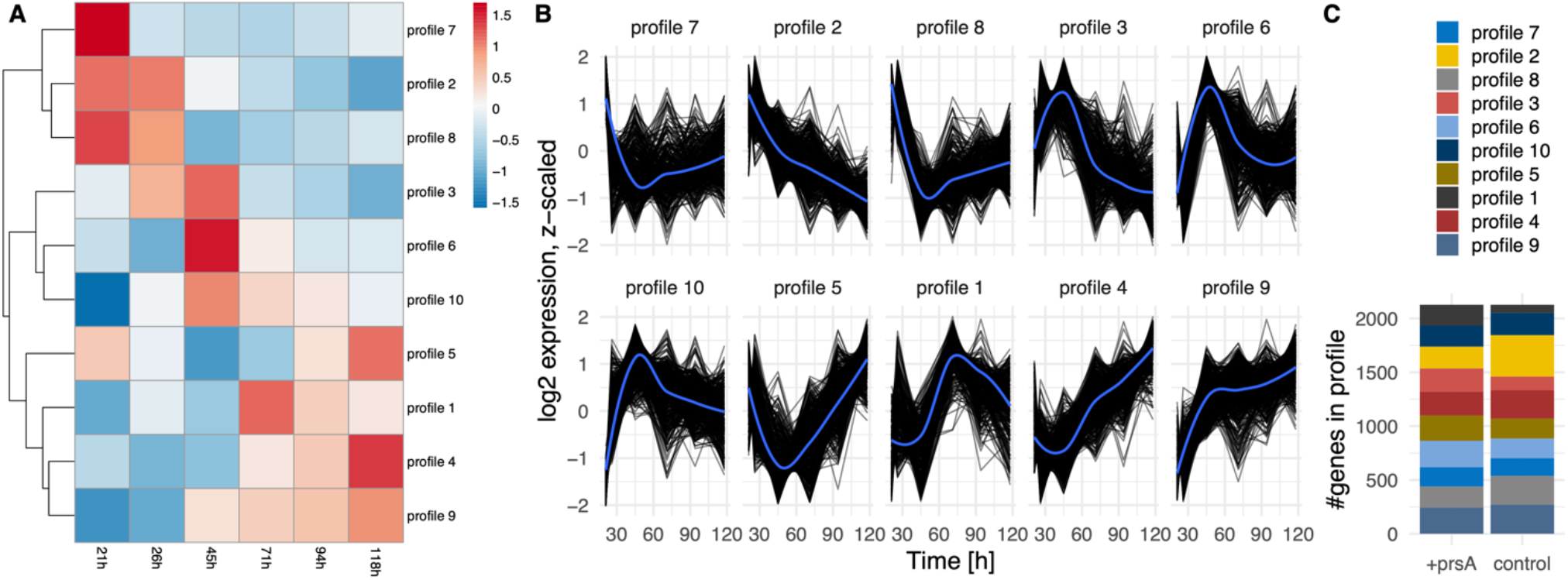
Expression profiles. **(A)** Heatmap of the expression profile over time (columns) for all differentially expressed coding and non-coding annotations investigated separately per strain. The resulting profiles were clustered (rows) and re-arranged by a complete linkage tree. **(B)** Profiles of expression per cluster for each annotation (black lines). An overall average expression according to a loess regression is added in blue. **(C)** The number of annotations per profile in either strain. The expression dynamics for each annotation can be in two separate profiles in the strains.

We assessed which biological processes (annotated in Gene Ontology, GO, terms (Ashburner et al., 2000)) are over-represented among the differentially expressed genes in each time and strain pairwise comparison (Fig 1 C). We compared the numbers of respective up- or downregulated genes relative to the number of expressed genes (see Methods). A total of 24 processes had significant over-representation (Fisher’s exact test, FDR p-adj. ≤ 0.01). We inspected the list of differentially expressed genes per process (Supplementary Table S7) in combination with meta-information available in the BSGatlas, particularly KEGG pathway annotations (Kanehisa and Goto, 2000; Geissler et al., 2021). Notably, the detected over-represented processes annotate genes with differentially expressed logFC predominately above the background logFC distribution of genes without detected differential expression (Figure S8). Further, some of the top 10 most extremely up- and downregulated genes (Table 2) were annotated by the detected processes (Table S7), namely cell wall macromolecule catabolic process (*safA* and *skfA*), response to stress (*nadC* and *nadE*), and ATP biosynthetic process (*ldh*). We further inspected the detected biological processes (Fig 3) for their relevance with respect to fed-batch fermentation, as described in the sections below.

**Figure 3.**
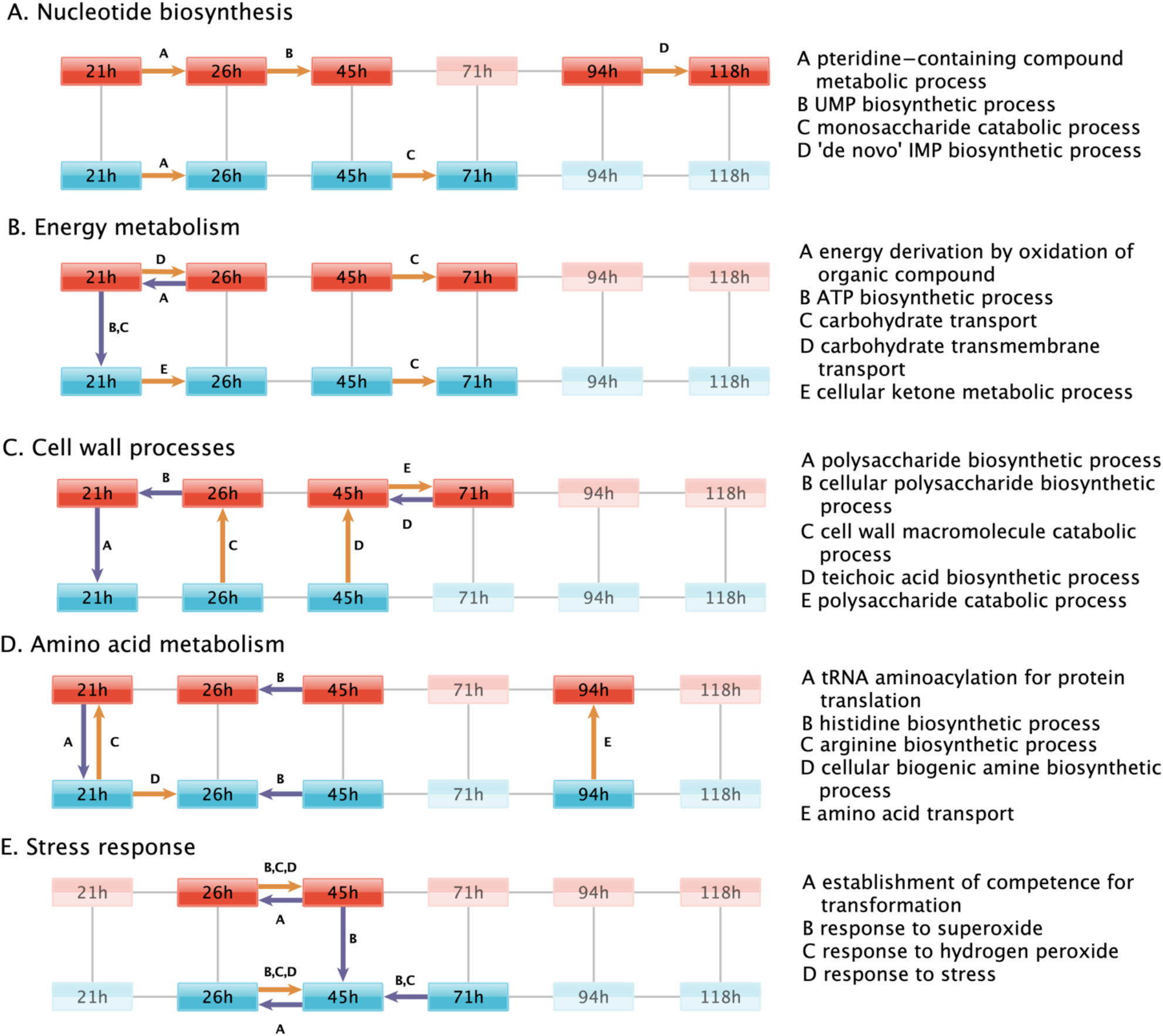
Regulated biological processes. Biological processes that are over-represented by the genes differentially expressed in each of the pairwise comparisons (black lines) between the fermentation timepoint in the +prsA (red) and control strain (blue). For simplicity, the regulated processes are grouped in subplots according to the same biological functions discussed in the result sections, which touch upon (A) nucleotide biosynthesis, (B) energy metabolism, (C) cell wall processes, (D) amino acid metabolism, and (E) stress response. Supplementary Fig S7 shows the regulated processes without further functional subdivision. Colored arrows indicate a pairwise comparison that was over-represented in a process (see description to the right). The arrows point to the conditions in which expression levels were higher. Upregulation in the +prsA strain or upregulation with time progression of the fermentation is highlighted in orange, whereas downregulation is shown in purple. In each subplot, time-strain conditions not adjacent to an arrow are greyed out.

#### 3.2.4 Nucleotide biosynthesis

It is well established that an ample supply of nucleotides is needed for efficient AMY protein expression (Hosoda et al., 1959), and thus also the nucleotide precursors, such as UMP and IMP, are of regulatory interest (Peifer et al., 2012; Hohmann et al., 2016b). Consistently, the over-representation investigation indicates an upregulation of UMP (GO:0006222) and IMP (GO:0006189) biosynthesis in the +prsA strain from timepoint 26h to 45h and 95h to 118h respectively. The monosaccharide catabolic genes (GO:0046365), especially the genes involved in the ribose synthesis via pentose phosphate pathway (Supplementary Table S7), are upregulated in the control strain from timepoint 45h to 71h. The pteridine-containing compound metabolic process (GO:0042558) was over-represented by genes upregulated from the first to the second timepoint in both strains. These specific genes are also part of the folate biosynthesis pathway, which is essential for both purine and pyrimidine synthesis (Kilstrup et al., 2005), and therefore quintessential for AMY production (Hohmann et al., 2016a; Hosseini et al., 2018).

### 3.3 PrsA over-expression affects genes involved in energy metabolism

#### 3.3.1 ATP biosynthesis

The ATP biosynthetic process (GO:0006754) was significantly downregulated in +prsA compared to the control strain on the first timepoint of the fermentation. Further, the data suggests that the energy derivation by oxidation of organic compounds (GO:0015980) was further downregulated in +prsA from the first to the second timepoint within the first day of fermentation. The differentially expressed genes associated with both processes comprise a long list (>50, see Supplementary Table S7) of core energy metabolic enzymes from the citrate cycle, oxidative phosphorylation, and glycolysis. Nevertheless, the list also overlaps with the starch and sucrose metabolism pathway, particularly with the glycogen biosynthesis (*glgA, glgB, glgC, glgD*, and *glgP*) (Kiel et al., 1994). Consistent with these observations, the carbohydrate transport (GO:0008643) was also downregulated in +prsA on the first timepoint. In contrast, the cellular ketone metabolic process (GO:0042180) was upregulated in the control strain from the first to the second timepoint. Ketones are essential for the biosynthesis of menaquinone (Lu et al., 2008). Menaquinone is *B. subtilis’* respiration coenzyme, similar in function to ubiquinone in human mitochondria (Lemma et al., 1990). Nevertheless, the ATP biosynthetic process (GO:0006754) was not detected significantly over-represented by the regulated genes at the other fermentation timepoints.

#### 3.3.2 Altering carbohydrate transport during fermentation

The over-representation analysis also suggests that both strains have an upregulated carbohydrate transport (GO:0008643, GO:0034219) from 45h to 71h. The transport might also be upregulated in the +prsA strain from the first to the second timepoint.

### 3.4 PrsA over-expression affects genes involved in cell wall destabilizing processes

Low PrsA protein abundances and increased concentrations of teichoic acid can reduce cell growth and cell wall disruption (Driessen et al., 1998; Hyyryläinen et al., 2000). For instance, the inhibition of the dlt operon–which is key to teichoic acid synthesis–increases AMY yields (Hyyryläinen et al., 2000; Yan and Wu, 2017). However, our data suggest that not only *dltB* expression is upregulaged in +prsA on timepoint 45h (logFC=0.86, adj. p<2.11e-5), but also the entire teichoic acid biosynthetic process (GO:0019350). Additional processes relating to cell wall molecules and polysaccharide biosynthetic (GO:0033692, GO:0000271) were observed as downregulated in +prsA. Nevertheless, not only does our data suggest that the biosynthesis is downregulated, the corresponding catabolic processes (GO:0016998, GO:0000272) might be upregulated.

## 4 Upregulation of amino acid metabolism during PrsA over-expression

### 4.1.1 Regulated amino acid metabolism

Genes of the arginine biosynthetic process (GO:0006526) are over-represented among the genes upregulated in the +prsA strain on the first timepoint and for the amino acid transport (GO:0006865) at timepoint 94h after fermentation started. The histidine biosynthetic process (GO:0000105) was detected as downregulated from timepoint 26h to the timepoint 45h in both strains. The data suggest also that the tRNA aminoacylation for protein translation (GO:0006418) is downregulated in +prsA on the first timepoint, and that the cellular biogenic amine biosynthetic process (GO:0042401) is upregulated in the control strain from the first to the second timepoint.

### 4.1.2 Expected changes in amino acid metabolism

Given the observed potential regulation in amino acid metabolism above, we investigated to which extend these might be the result of the peptide sequence of the secreted AMY. The inspection of codon composition of all coding genes suggests that the AMY and the over-expressed PrsA contain substantially more tryptophan, asparagine, aspartic acid, and lysine (more than 2 standard deviations from the average proportion, Table 3). Tryptophan was the strongest over-represented amino acid in AMY (+3.1 standard deviations). But in comparison, the subset of differentially expressed coding genes did not change the overall composition (within 1 standard deviations). Given the high energetic cost of tryptophan biosynthesis (Akashi and Gojobori, 2002), the evolutionary adapted amino acid metabolism will be affected (Smith and Chapman, 2010).

**Table 3.**
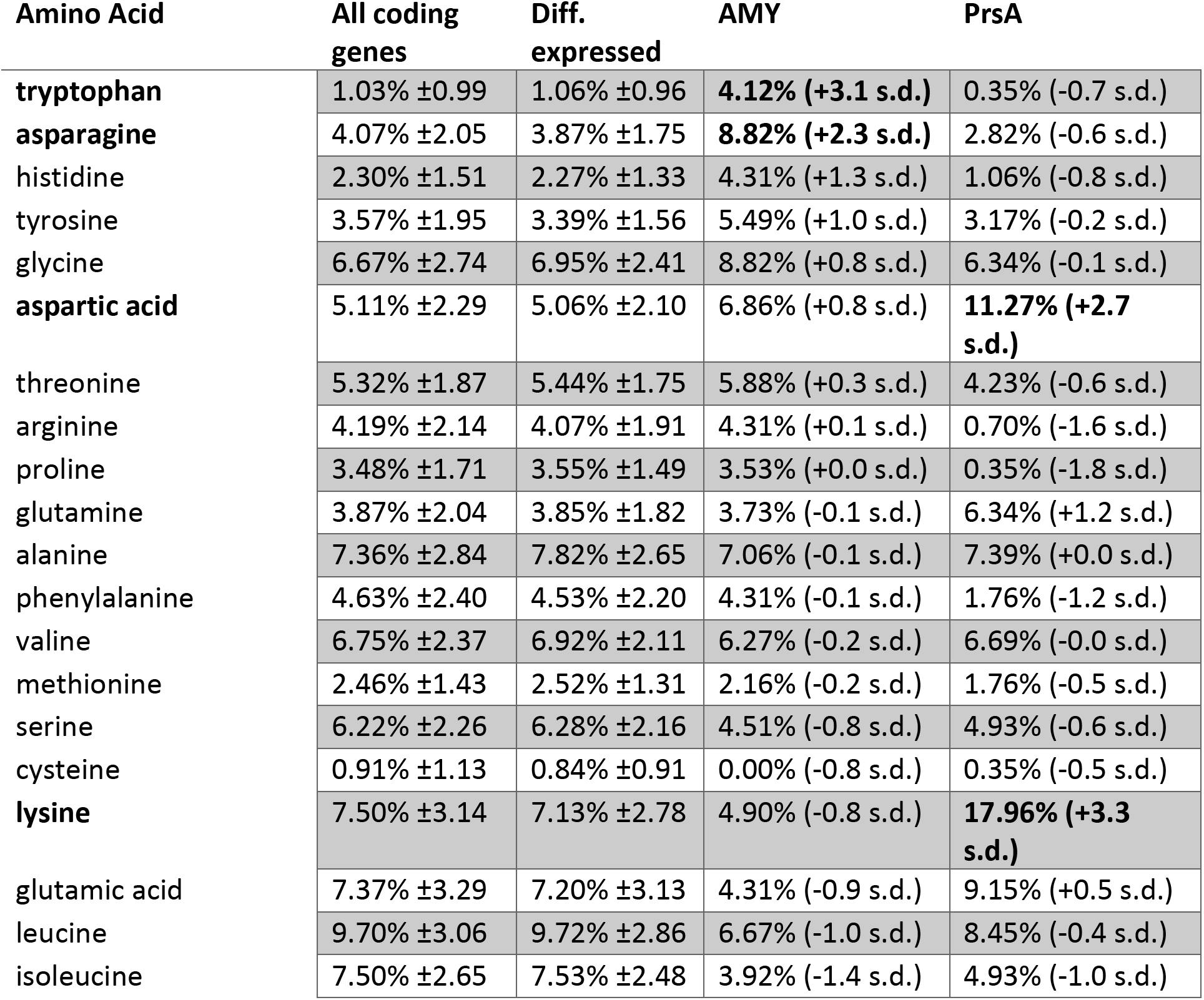
Amino acid composition. The average amino acid compositions (rows) are shown for all coding genes (second column) and those that were detected as differentially expressed (third column). The standard deviations are shown behind the “±” signs. The compositions of amino acids for the AMY enzyme (fourth) and the over-expressed PrsA (fifth) column are shown. The difference in standard deviations relative to the average for all genes are indicated in parenthesis. The bold font highlights amino acids with difference of more than 2 standard deviations.

### 4.2 Protein-protein interactions of stress response and competence transformation

#### 4.2.1 Stress response turning point

The over-representation investigation reveals that both strains upregulate parts of their stress response concerning the reactive oxygen species (ROS) response (GO:0006950 and the two children terms GO:0042542, GO:0000303) from timepoint 26h to 45h. Simultaneously, the strains downregulate the establishment of competence for transformation (GO:0030420). The protein ClpC is the key switch between heat shock (including secretion stress) and competence regulation (Turgay et al., 1997). During stress, a three protein complex of ClpC, MecA, and ComK is formed (Turgay et al., 1997). The bound central competence regulator ComK can no longer act as a transcription regulator, which prevents the establishment of competence (Turgay et al., 1997). According to our results, *clpC* undergoes significant differential expression during fermentation in both strains, but neither *comK* nor *mecA* had significant expression changes though both were expressed (Supplementary Table S4). Given that the molecular mechanism of the ClpC switch (i) is post-translational, (ii) does not directly impact the transcription levels of the involved genes, and (iii) involves a third factor, the analysis by pairwise comparison of expression levels cannot detect that specific interaction. Therefore, we complemented the expression analysis with a protein-protein interaction (PPI) network analysis.

#### 4.2.2 PPI network analysis

We retrieved PPIs from the STRING database for the *B. subtilis* strain 168. STRING provides a list of functional associations from multiple evidence channels, such as curated knowledge from known metabolic pathways and protein complexes, physical PPIs from lab experiments (*e.g*., pull-down assays), predicted interactions from text mining of the biomedical literature, or associations based on co-expression analysis (Szklarczyk et al., 2019). The resulting network of 4,774 high-confidence associations (confidence score >0.8) among 1,770 of the 1,791 differentially expressed protein-coding sequences was clustered into 201 protein clusters using MCL (Enright, 2002; Morris et al., 2011; Doncheva et al., 2019). In combination with the significant logFCs between the +prsA and control strains (Legeay et al., 2020), we manually inspected 4 clusters with interesting patterns regarding this study’s outset (Fig 4). These are described in the following sections below.

**Figure 4.**
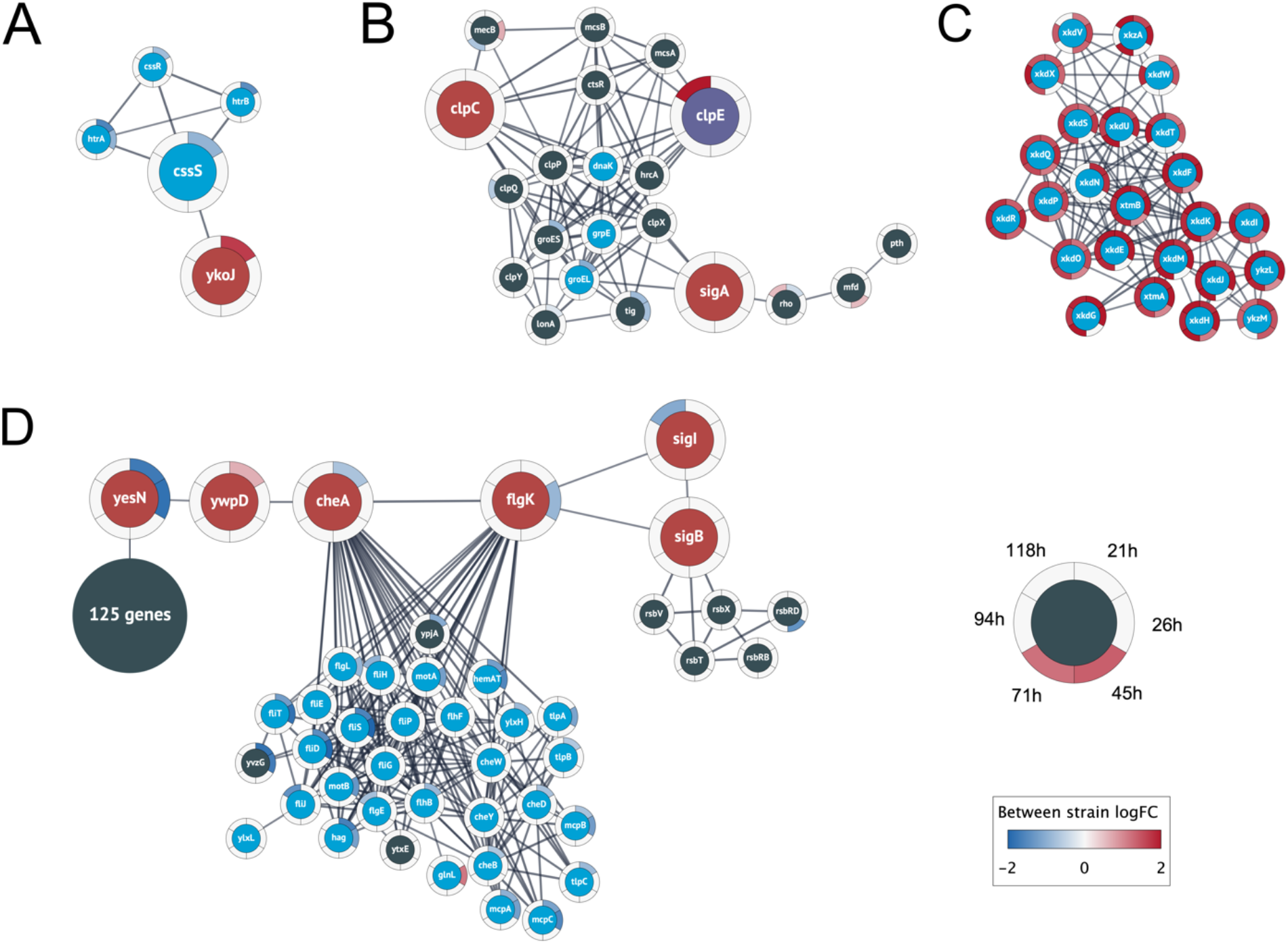
Relevant clusters of differentially expressed genes. Nodes represent protein coding genes and edges correspond to high-confidence protein interactions retrieved from STRING. The differential expression between strains is shown as rings around the nodes, where each ring contains the logFC values for each time point comparison in a blue-white-red color gradient (see figure legend). A high positive logFC is colored red and indicates a significantly larger expression in the +prsA strain compared to the control. Non-significant differential expression is shown as 0 logFC (white). The logFC color gradient was truncated at ±2. (A) The genes in this cluster include the central heat shock stress two-component system of CssRS and the proteases HtrAB (blue nodes). The cluster also contains the gene ykoJ of unknown function (red node) connected to the stress transducer CssS (large blue node). **(B)** This cluster contains the competence/heat shock switch protein ClpC (leftmost red node) and the universal sigma factor SigA (rightmost red node); SigA and ClpC share interactions with the tree heat shock proteins dnaK, grpE, and groEL (blue nodes). The cluster also contains ClpE (purple node) that had substantially higher expression in +prsA at timepoint 118 h (logFC ~2.6). **(C)** The analysis found a cluster of 24 prophage or prophage-like genes that were closely interacting and had significantly higher expression in +prsA throughout the fermentation. **(D)** The largest cluster contains a “bottleneck” of high-confidence interactions at two genes of unknown function (yesN and ywqD) between 125 genes of various catalytic function (summarized as one node) and 29 chemotaxis genes (blue nodes) and the central chemotaxis signal protein CheA, the flagellar hook-filament flgK, the general stress repose sigma factor SigB, and the RNA polymerase sigma factor SigI.

#### 4.2.3 Two-component system

The first PPI cluster consists of the CssRS two-component system, including the involved proteases (see Introduction, Fig 4 A). However, the cluster contains an additional association between the stress signal transducer CssS and YkoJ of unknown function. The *ykoJ* expression during secretion of a vaccine compound (beta-toxoid) positively depends on CssS (Nijland et al., 2007). In contrast, the expression during AMY might have a negative dependency with *cssS* being significantly lower expressed in +prsA on timepoint 21h after fermentation start (logFC = −0.9, adjusted p = 8.5e-10) and *ykoJ* significantly higher (logFC = 1.7, adj. p = 1.2e-7). To our knowledge, the association YkoJ-CssS has not been characterized in the context of AMY production.

#### 4.2.4 Competence switch

The second cluster (Fig 4 B) contains the above-described heat shock/competence protein switch ClpC (Turgay et al., 1997). The cluster also contains ClpC’ repressor CtsR (Derré et al., 1999) and the universal sigma factor SigA. Further, SigA and ClpC share associations with the three heat shock proteins DnaK, RrpE, and GroEL. Although *mecA* was not detected as differentially expressed, the paralog *mecB* was, and it is part of this second cluster (Persuh et al., 2002). *B. subtilis’* other two Clp-proteins ClpP and ClpE are also part of this cluster. ClpE had a significantly higher expression on timepoint 118h in +prsA (logFC = 2.6, adj. p = 0.0005), which is relevant because ClpE destabilizes the functionality of the repressor CtsR (Miethke et al., 2006).

#### 4.2.5 Prophage genes

A third cluster (Fig 4 C) contains a set of tightly associated 24 PBSX prophage and prophage-like genes that were all significantly higher expressed in +prsA compared to control at various timepoints during the entire duration of the fermentation. PBSX, a defective *B. subtilis* prophage (Wood et al., 1990a), is known to be potentially heat-induced (Wood et al., 1990b), and they have a potential association with the level of lytic stress resistance (Buxton, 1980).

#### 4.2.6 Potential cell motility regulation

Finally, the fourth cluster has an interesting pattern of associations involving many chemotaxis genes (Fig 4 D). This cluster is structured into two separate interconnected components: On the one side there are 29 chemotaxis proteins and on the other 125 protein-coding genes with various catalytic functions (116 of 125 [92.8%] genes are annotated in the general catalytic activity term GO:0003824), however, both parts are connected by a backbone of associated genes. This backbone includes the central flagella motion frequency regulator CheA, the flagellar hook-filament FlgK, the general stress sigma factor SigB, the heat-shock protein sigma factor SigI, and the two partially characterized signal transducers YesN and YwspD (Fabret et al., 1999; Petersohn et al., 2001; Zuber et al., 2001; Asai et al., 2007; Mukherjee and Kearns, 2014). Interacting with sigB are 5 stress regulatory proteins induced by SigB (according to STRING annotations). Both YesN and YwsqD are described as histidine kinases, although the corresponding response regulator remains unknown (Fabret et al., 1999; Caspi et al., 2014; Zhu and Stülke, 2018; Geissler et al., 2021). Even if the regulators are unknown, the backbone has an interesting pattern of antagonistic logFC: (i) YesN is significantly lower expressed in +prsA on timepoint 21h (logFC=-1.7, adj. p=1.4e-6) and 26h (logFC=-1.84, adj. p=8.5e-5), (ii) YwpD is higher expressed in +prsA on 21h (logFC=0.6, adj. p=0.0037), and (iii) CheA lower again on 21 h (logFC=-0.7, adj. p=0.0025). The bottom-line is that the PPI analysis elucidates the tight associations between heat shock, competence transformation, cell motility, general stress response, and translation.

## 5 Discussion

In this study, we investigated how PrsA over-expression in *B. subtilis* impacts the transcriptome during fed-batch alpha-amylase (AMY) fermentation. We carried out a temporally resolved RNA-seq study to analyze expression levels and regulation of biological processes with respect to secretion stress. We inspected a comprehensive set of coding and non-coding reference annotations as well as 542 novel potentially transcribed regions (NPTRs). The fermentation process strongly affect gene expression and we observe a large number of differentially expressed genes both between the strange and overtime: At total of 1,793 coding genes (67% of expressed genes), 234 NPTRs (66%), 68 putative ncRNAs (64%), 20 riboswitches (54%), 9 tRNAs (41%), and 3 sRNAs (33%) were differentially expressed. The PrsA over-expressing strain, which is consistent with prior descriptions had increased yield and reduced growth (Quesada-Ganuza et al., 2019), was observed to have significant strain-specific differential expression for more than half of the transcribed genes. Subsequent in-depth analysis of regulated biological processes (Fig 3) and the PPI network of differentially expressed coding genes (Fig 4) shed light on the complex intertwined processes of stress pathways, the core energy metabolism, and cell motility (Helmann et al., 1988; Marquez et al., 1990; Storz and Hengge, 2010; Yan and Wu, 2019).

### 5.1 Amino acid and energy metabolism

The observation of the potentially downregulated ATP biosynthesis in the +prsA strain surprised us: (i) The AMY hypersecretion is stressful and energy-intensive for the cells (Song et al., 2015). (ii) It has been hypothesized that ATP might be required for PrsA chaperone activity (Yan and Wu, 2017). (iii) The reduction of ATP levels can also increase the general stress response of *B. subtilis* (Haldenwang, 1995; Petersohn et al., 2001; Yan and Wu, 2019). The potential downregulation of ATP biosynthesis in the +prsA strain seems counterintuitive because the strain has both lower stress and higher yield than the control (Quesada-Ganuza et al., 2019). However, the reduced ATP biosynthesis might be due to the impact of the hypersecreted AMY and over-expressed PrsA on the amino acid metabolism. Contrary to the evolutionary energetic adaption of the amino acid composition for secreted proteins (Smith and Chapman, 2010), the four amino acids tryptophan, asparagine, aspartic acid, and lysine are over-represented in the AMY and PrsA proteins (Table 3). Although, the specific metabolism processes for these four amino acids were not detected as significantly regulated during fermentation (Fig 3), more general amino acid processes (*e.g*., transport) or biosynthetic processes for other amino acids (arginine and histidine) were significantly over-represented by regulated genes. On the one hand, the upregulation of arginine synthesis and related transport mechanisms improves osmotic stress resistance (Du et al., 2011; Zaprasis et al., 2015), which in turn is beneficial to AMY production in *B. subtilis* (Zhao et al., 2018). On the other hand, the over-represented amino acids might explain the reduced ATP biosynthesis. (i) Tryptophan is the amino acid with the highest biosynthetic cost in *B. subtilis*, with a 42.9% higher cost than the second most costly amino acid (phenylalanine) (Akashi and Gojobori, 2002).(ii) The biosynthesis, in particular for costly amino acids, diverges intermediate metabolites from ATP biosynthesis (Akashi and Gojobori, 2002). In the case of tryptophan, the intermediate metabolites are already diverged from glycolysis, which also impacts the downstream citrate cycle (Kanehisa and Goto, 2000; Akashi and Gojobori, 2002). However, a more definite inspection to confirm the regulation of the amino acids and ATP metabolism would require investigation of concentrations of the individual metabolites with for instance metabolomics.

### 5.2 Cell wall destabilizing processes

The over-expression of PrsA is known to lead to reduced cell growth and cell lysis (Quesada-Ganuza et al., 2019). It was suggested that protein-protein interactions of specific PrsA protein domains are causal for these phenotypes (Quesada-Ganuza et al., 2019). Our data suggest that, on a transcription regulatory level, the PrsA-over-expressing stain has both increased polysaccharide catabolism and reduced polysaccharide biosynthesis. We hypothesize that this strongly contributes to cell wall breakdown, which leads to the detrimental phenotypes. Therefore, investigating the associated differentially expressed genes could potentially be the outset to trace back the causality chain of why their regulation changes, and as path forward to finding candidates that stabilize cells walls and increase yields. Further, the PPI network analysis highlighted 24 tightly-associated PBSX prophage and prophage-like genes (Fig 6 C) that might be decisive in unraveling the PrsA over-expression lysis phenomena (Buxton, 1980; Quesada-Ganuza et al., 2019), particularly due to the heat-induced (and thus secretion stress related) expression of the PBSX genes (Wood et al., 1990b).

### 5.3 Stress and cell motility

The PPI network analysis resulted in four clusters of proteins that we found to be relevant to this study’s outset (Fig 4). These were the genes of the CssRS two-component secretion stress response in one cluster (Fig 4 A), while the known ClpC regulatory switch and its associations with secretion stress, competence transformation, and associations with the universal sigma factor SigA belong to another cluster (Fig 3 B) (Turgay et al., 1997). Further, the analysis provided a large cluster (Fig 4 D) of cell motility-related genes, which is consistent with the large number of proteins involved in regulating bacterial motility (Rajagopala et al., 2007). The closer inspection of the latter cluster suggests that the proteins YesN and YwsqD might have a signaling role in balancing between cell motility and 125 genes that are annotated to have various metabolic catalytic functions*, e.g*. the phosphogluconate dehydrogenase. To our knowledge, the potential relationship between cell motility and AMY fermentation has not been elucidated so far, although a potential hypothesis could be that the signaling facilitates the regulation of flagellar cell motility to escape from the stress region (Helmann et al., 1988; Marquez et al., 1990; Yan and Wu, 2019). Nevertheless, a follow-up study is needed to verify cell motility regulation during AMY production.

### 5.4 Conclusion

In conclusion, our transcriptome study highlights the expression dynamics of secretion stress during fed-batch AMY fermentation. The comparison of expression levels in a PrsA over-expressing strain to a control strain showed differential expression for nearly half of the transcribed genes. A wide variety of up- and downregulated biological processes related to energy and amino acid metabolism. Also, the data shows potential associations of the cell lysis phenomenon of PrsA over-expression with the stress response and cell motility. Overall, these results identify genes and biological processes, which are affected during fermentation and by the overexpression of PrsA and provides a starting point for future genetic modification of B. subtilis for improved yield.

## Supporting information

Supplementary Sequences

## 6 Data Availability

The genomic sequences and RNA-seq data were deposited in the GEO database (GSE189556). The expression coverages are presented as a browser for interactive investigation at (https://rth.dk/resources/prsa/). The annotations of the BSGatlas are accessible at https://rth.dk/resources/bsgatlas/. The additional putative ncRNA annotations are part of the supplementary information of (Nicolas et al., 2012). The RNA-seq data was processed with a reproducible pipeline located at doi 10.5281/zenodo.4534403.

## 7 Acknowledgments

This work was supported by the Innovation Fund Denmark [5163-00010B] and the Novo Nordisk Foundation [NNF14CC0001]. The authors thank Annaleigh Ohrt Fehler for pivoting the samples for sequencing and Thomas B Kallehauge for support in conducting the fermentations and sampling.

## 8 Author Contributions

ASG conducted the entire computational analysis and wrote the manuscript. LDP extracted the RNA. NTD contributed to the analysis and methodology design of the PPI network. CA contributed to the discussion of the expression analysis. EGT contributed to the swriting in the early-stage. AB prepared the bacterial strains. LJJ contributed to discussion of the gene clustering, enrichment analysis, and PPI network analysis. SES, CH, JV, JG supervised the work. JG and ASG made the study design. JG was the main project coordinator. All authors read and approved the manuscript.

## 9 Contribution to the Field Statement

Our manuscript provides an in-depth RNA-seq study into the expression dynamics of secretion stress during fed-batch amylase fermentation. The analysis of differentially expression shows regulation for nearly half of the *Bacillus subtilis* transcriptome. We found many up- and downregulated biological processes ranging from the energy and amino acid metabolism to cell wall synthesis. A complementary protein association network analysis sheds light on the potential associations between the stress response and cell motility in the context of PrsA over-expression. Overall, these results form the basis and outset for future study in the field of yield optimization.

## 10 Conflict of Interest

The authors declare that the research was conducted in the absence of any commercial or financial relationships that could be construed as a potential conflict of interest.

## Notes

### Competing Interest Statement

The authors have declared no competing interest.

