## Supplementary Figures for "The impact of PrsA over-expression on the *Bacillus subtilis* transcriptome during fed-batch fermentation of alpha-amylase production"

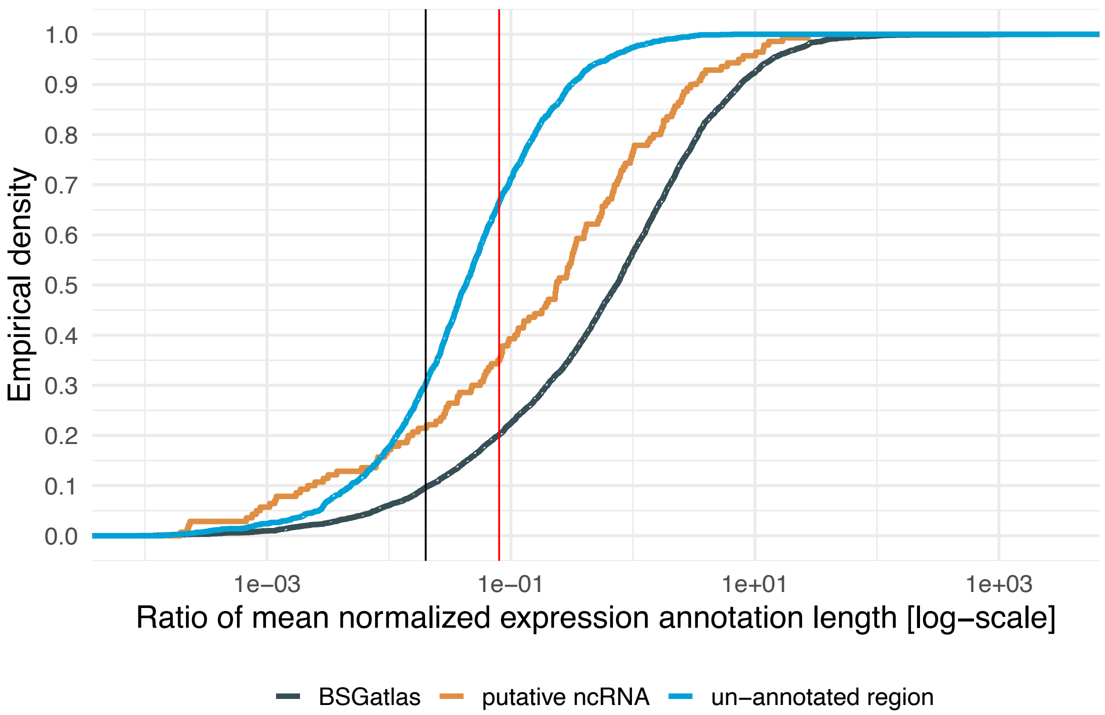


**Figure S1. Expression in Un-Annotated Regions** Gaps between annotations, the un-annotated regions, might still harbor valuable expression signals. In order to select a minimal expression level cutoff, the ratios of DESeq2 normalized expression relative to the length were inspected for the known annotations. The un-annotated regions (blue, listed in Table S3) were inferred when considering available annotations from the BSGatlas (black) and the additional putative ncRNA (orange, see manuscript). The figure shows the cumulative empirical density for the ratios. With the 50 bp read lengths in mind, the black vertical line indicates a one-fold coverage and the red line a four-fold coverage of an annotation.

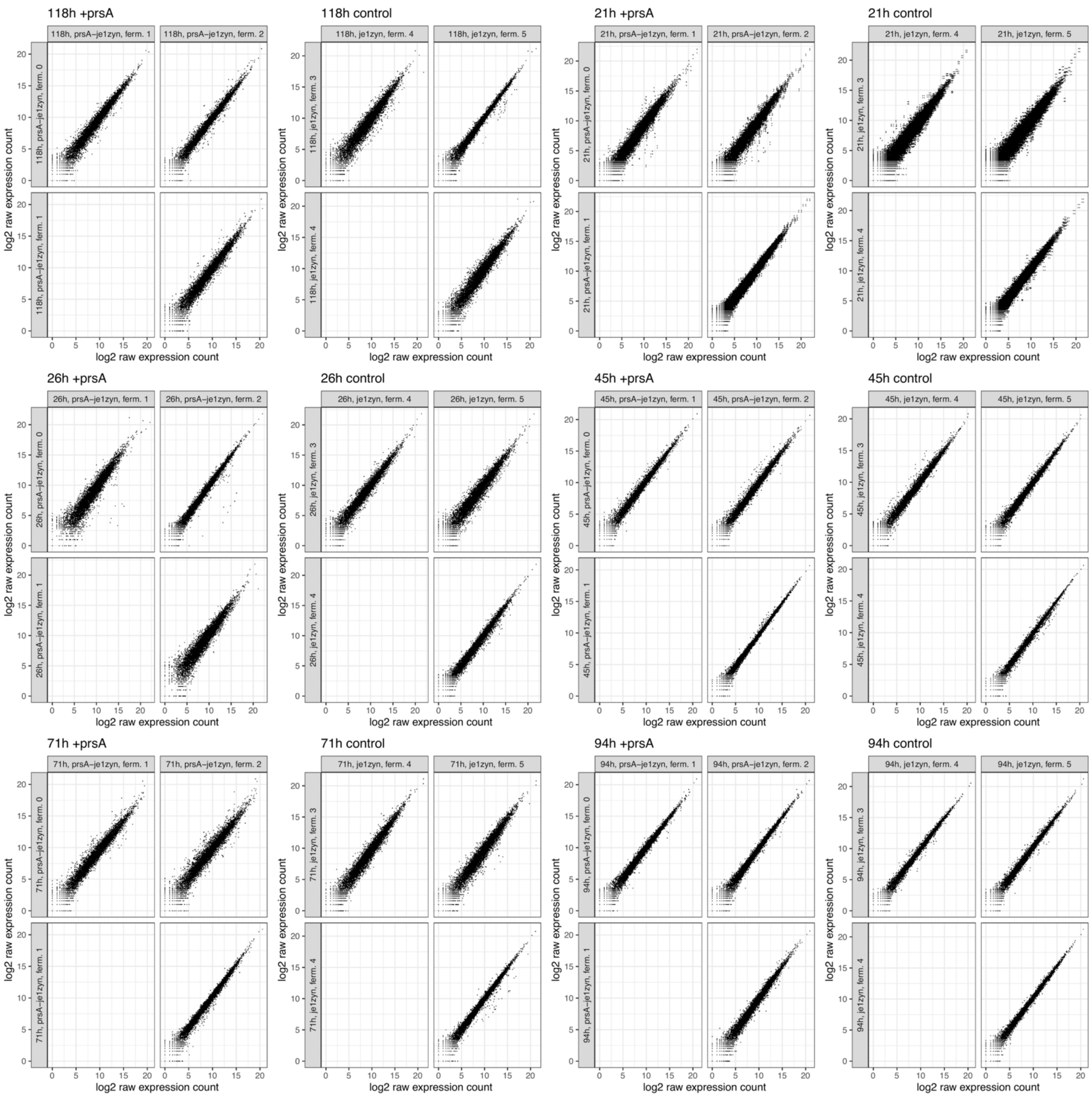


**Figure S2. Expression Scatterplot.** For each timepoint and strain condition, shown are the scatterplots between the triplicate samples in a timepoint+strain condition (sub-panels). The biological replicates are numbered ferm. 1 through 6 (columns and rows). The scatterplots are relative to the raw log2-transformed expression counts without further normalization.

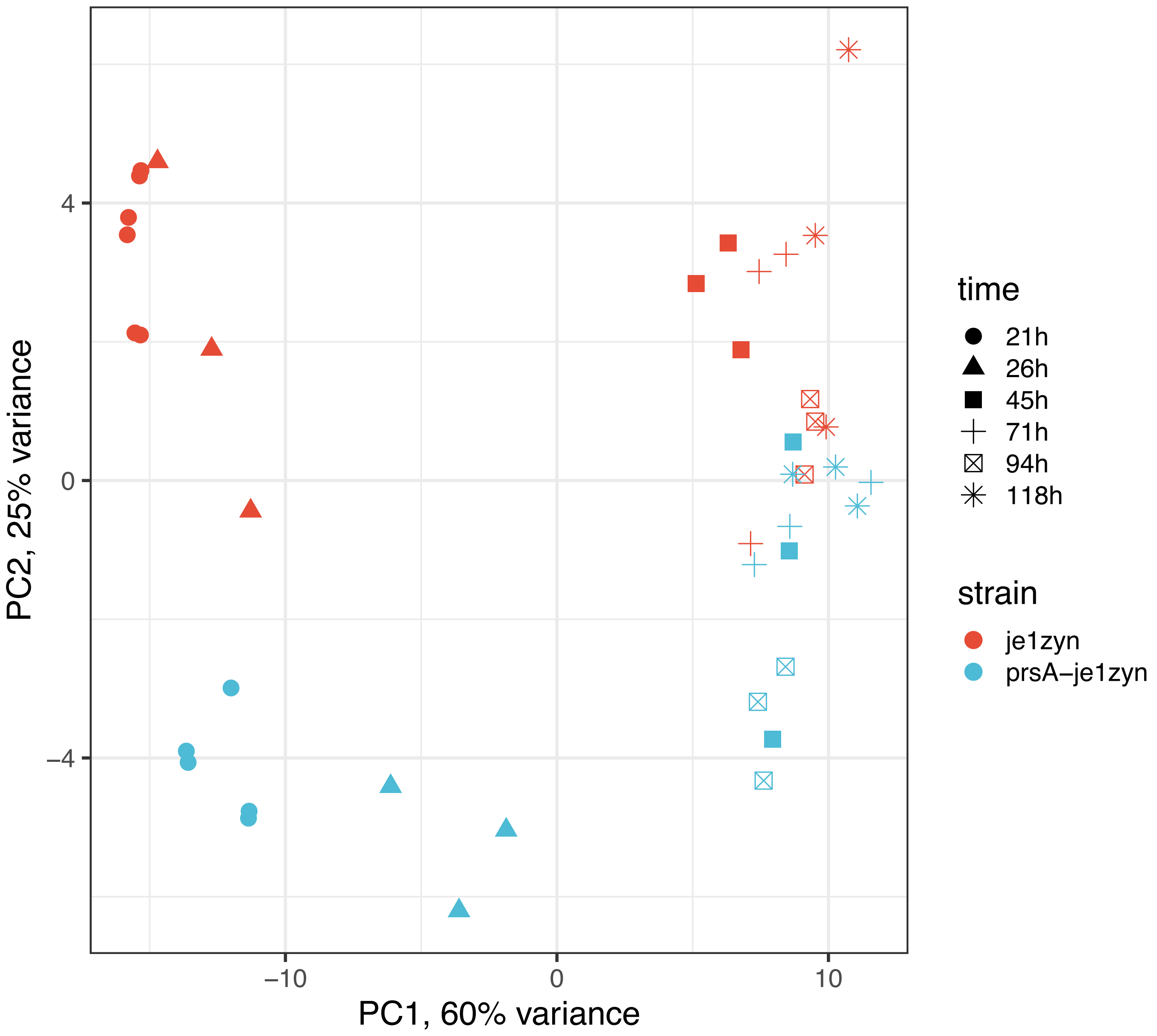


**Figure S3. PCA of libraries.** The PCA shows the RNA-seq libraries for the top 100 genes by expression variance after blind regularized log transformation (DESeq2). No differential expression was conducted for this figure. The shape indicates the timepoint at which a sample was taken (see legend) and the color distinguishes the control strain (red) and the PrsA over-expressed strain (blue).

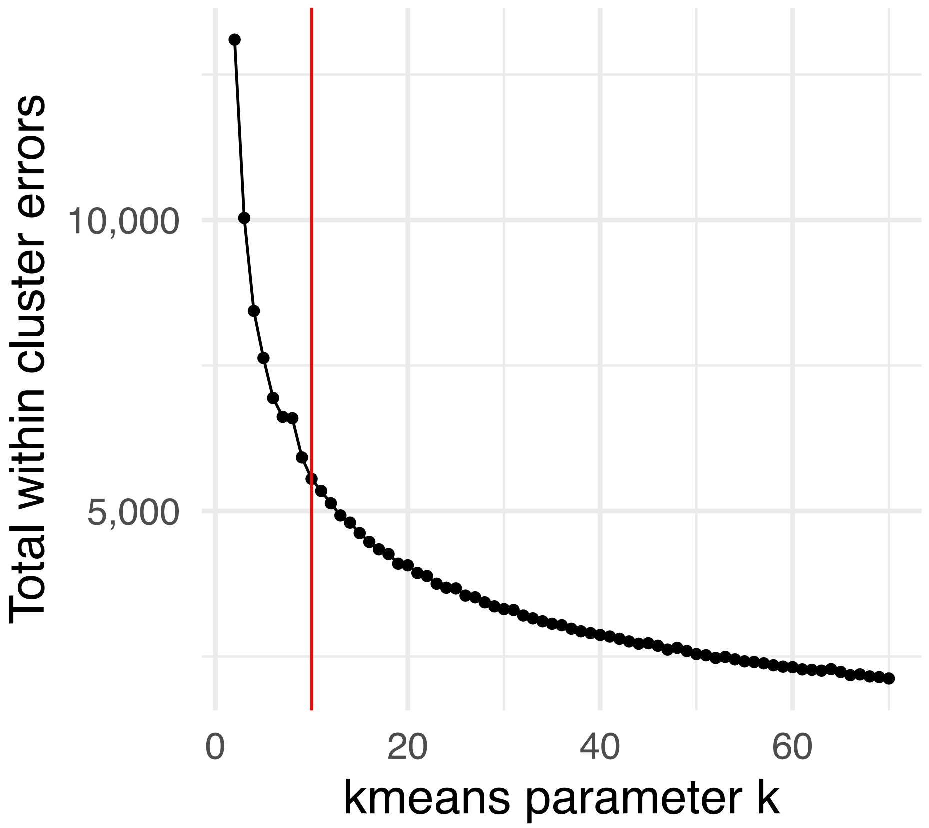


**Figure S4. K-means Clustering.** After differential expression analysis, genes with similar expression levels (according to their log expected expression, Table S5) were clustered together with k-means. The parameter was selected (red line) according to the elbow plot which shows the total within-cluster errors for all possible k=2…70.

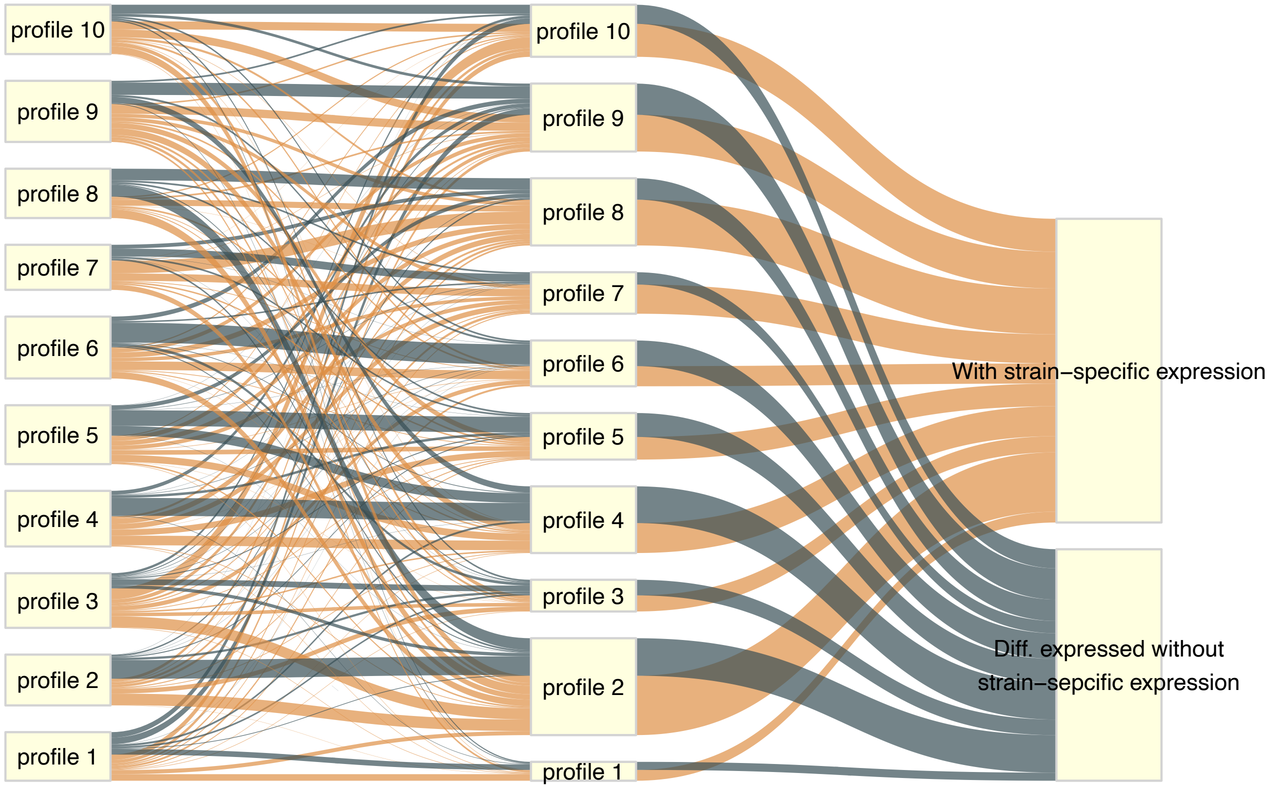


**Figure S5. Sankey.** The differential expression analysis revealed which genes had strain-specific expression patterns (Table 2). Based on the mean expression levels clustering (Figure 2), this Sankey plot shows the expression profiles that the differentially expressed genes with strain-specific expression patterns (orange) and those without (blue) had.

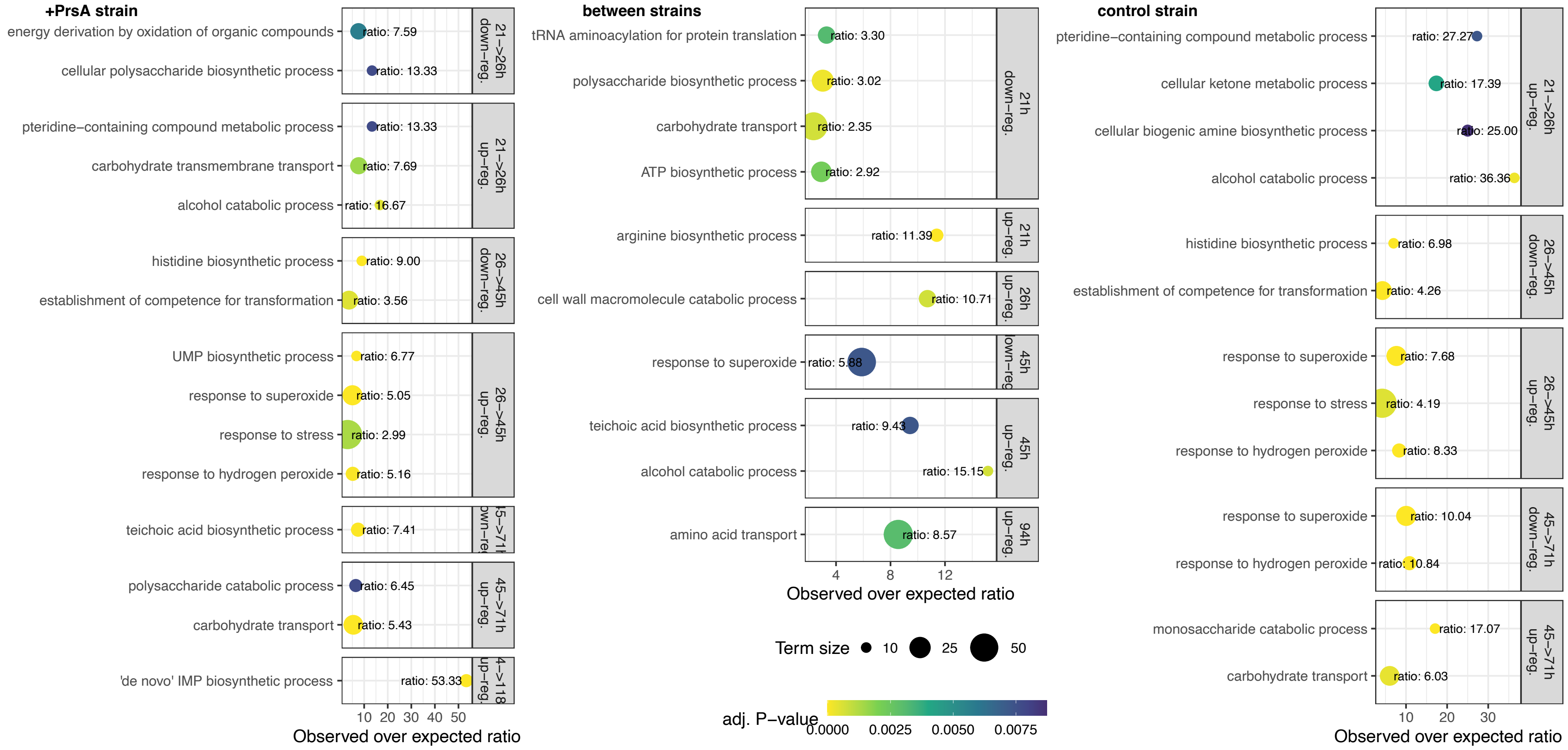


**Figure S6. Over-represented biological processes.** For each pairwise differential expression analysis (Fig 1 C), the regulated biological processes were detected via over-representation analysis (see methods). The over-represented regulated processes during fermentation of the +prsA strain (left), the control strain (right), and those with differences between the strains (middle) are shown per test with a point that scales with the background size of the process (see methods). The color corresponds to the FDR adjusted P-value of over-representation. The x-axis is the ratio of observed regulated genes to those expected by chance (term size over the number of up- or down-regulated genes in pairwise test).

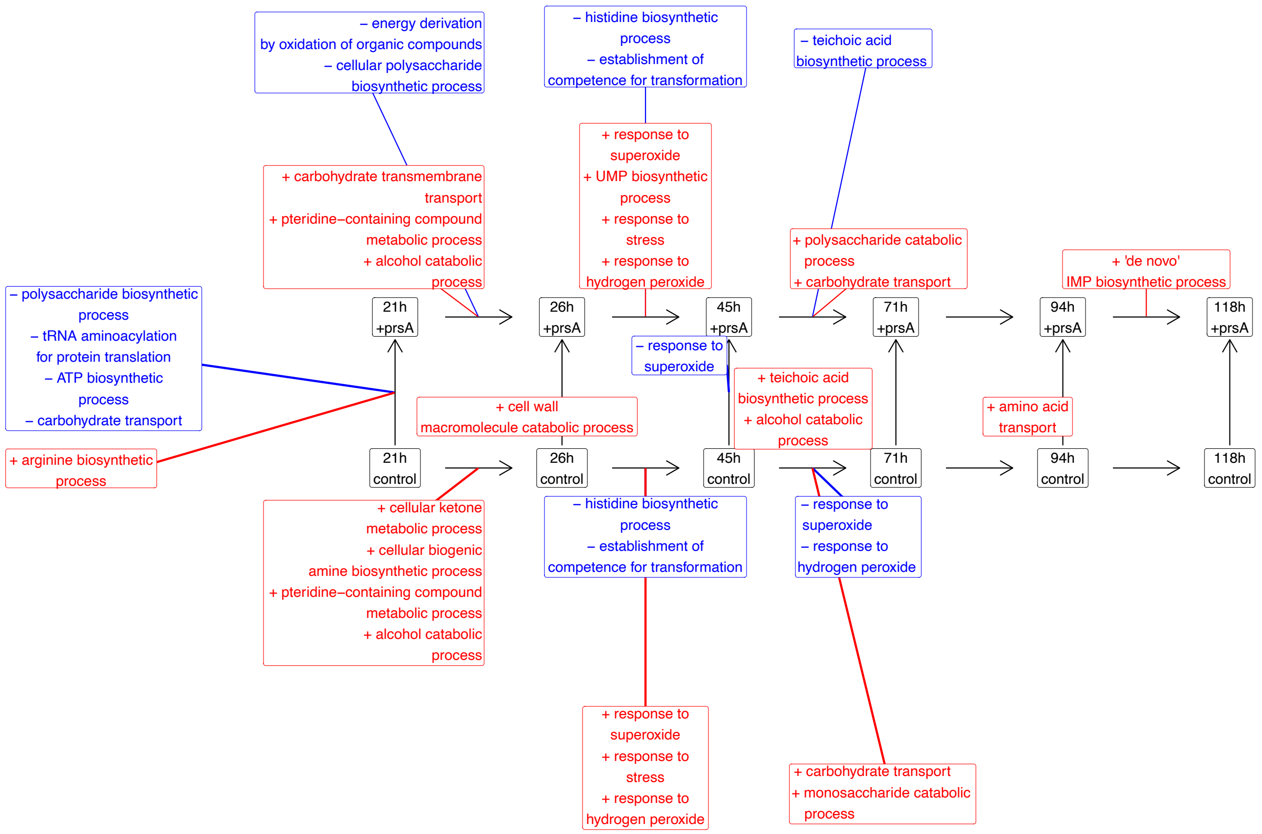


**Figure S7. Enriched biological processes.** Shown are the GO biological processes that were over-represented by either all up- (red, plus sign) or down-regulated (blue, minus sign) genes differentially expressed in one of the pairwise comparisons (arrows). The up- and down-regulations are in the direction of the arrows, *e.g.*, the “ATP biosynthetic process” was over-represented by gene upregulated in the +prsA strain at the first timepoint.

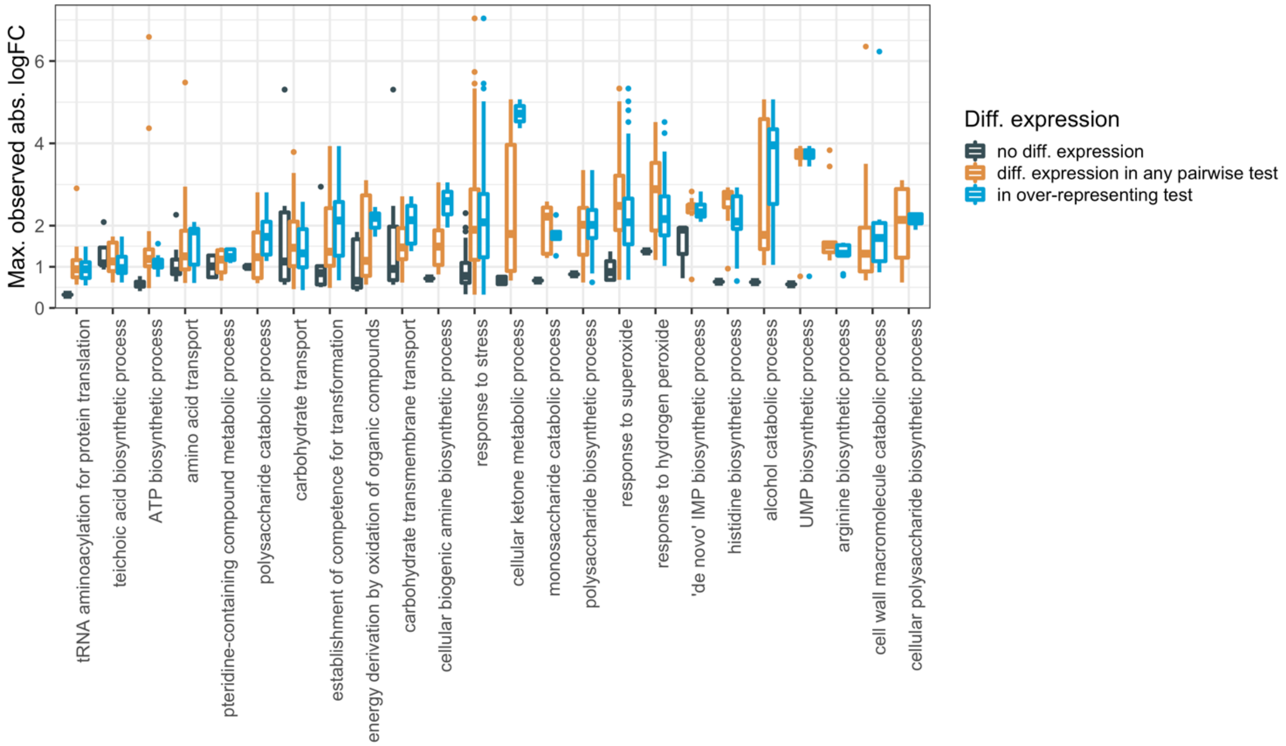


**Figure S8. Comparison of logFC in over-represented processes.** For each of the over-represented biological processes from Figure S7 (x-axis), the boxplots shows the distribution of absolute logFCs (y-axis) for genes that are part of the respective processes in three different cases: In the first case, the logFCs relates to those observed in the pairwise test which was over-represented in the respective process (blue). Second, the distribution relates to the maximal differentially expressed logFC for any pairwise test (orange). Lastly, the shown logFCs are the maximum observed for genes for which differential expression was not detected (black).

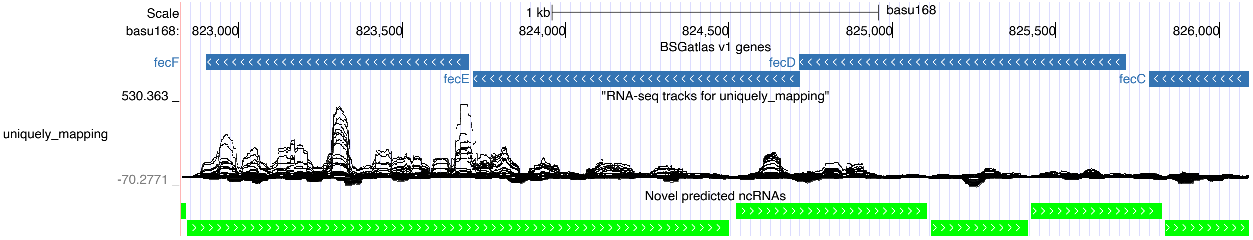


**Figure S9. Fragmentation of read coverage based-ncRNA prediction.** Shown is a genome browser snapshot of the region S257, which is anti-sense located to the fecCDEF genes (blue). The normalized RNA-seq coverage signal of this study is shown underneath the genes. At the bottom, green bars indicate predicted ncRNA according to a pipeline that uses the coverage signal (Geissler et al). Due to drastic drops in the coverage signals, the S257 region is covered by 6 distinct predicted regions, although this region has been described to be transcribed as one (Nicolas et al, 2012).
