## Supplementary Sequences for "The impact of PrsA over-expression on the *Bacillus subtilis* transcriptome during fed-batch fermentation of alpha-amylase production"

>Synthetic AMY JE1

ATGAAACAACAAAAACGGCTTTACGCCCGATTGCTGACGCTGTTATTTGC

GCTCATCTTCTTGCTGCCTCATTCTGCAGCCGCGGCACATCACAATGGTA

CGAATGGGACGATGATGCAGTATTTCGAGTGGCATCTGCCGAATGACGGC

AACCATTGGAACAGGCTGCGGGACGACGCGTCCAATTTAAGGAATCGCGG

CATAACTGCGATCTGGATACCACCGGCGTGGAAGGGTACGTCACAAAACG

ACGTGGGATATGGAGCGTACGACCTCTACGATTTAGGAGAATTCAACCAG

AAAGGCACAGTTCGAACAAAGTACGGGACGCGCTCTCAATTGGAGTCAGC

GATCCATGCTCTGAAAAACAACGGCGTTCAAGTTTATGGCGATGTTGTCA

TGAACCATAAAGGCGGTGCAGATGCGACAGAAAATGTGCTCGCAGTTGAA

GTAAATCCAAACAATCGAAACCAAGAGATTTCCGGAGACTATACGATCGA

AGCCTGGACAAAATTTGATTTTCCAGGTCGGGGCAACACATATAGTGATT

TTAAATGGCGCTGGTACCATTTCGACGGCGTAGATTGGGATCAGTCACGG

CAATTCCAAAACCGAATCTATAAGTTTAGAGGCAAAGCTTGGGACTGGGA

AGTTGATTCGGAAAACGGAAACTATGATTATTTAATGTATGCGGACGTGG

ATATGGATCATCCGGAGGTGGTGAACGAACTTCGCCGTTGGGGGGAATGG

TATACCAATACGTTAAACCTGGATGGATTTCGCATAGATGCCGTGAAACA

TATTAAATACTCATTTACCCGGGATTGGTTAACGCATGTCCGGAACGCCA

CGGGTAAAGAGATGTTCGCCGTCGCGGAATTTTGGAAAAATGACCTGGGC

GCCTTGGAAAATTACCTTAACAAAACGAACTGGAATCACAGCGTCTTTGA

CGTACCGCTTCACTACAACTTATATAATGCATCTAACTCAGGAGGCAATT

ATGACATGGCCAAGCTTTTGAATGGAACGGTGGTTCAAAAGCACCCGATG

CATGCAGTGACGTTCGTCGATAACCATGATTCACAGCCTGGGGAGTCCCT

CGAAAGTTTCGTCCAGGAGTGGTTCAAACCATTAGCATATGCCCTTATTT

TGACGAGGGAACAAGGATATCCTAGTGTTTTTTACGGCGACTATTATGGA

ATCCCGACACATTCTGTGCCGGCCATGAAGGCAAAAATCGATCCAATCTT

GGAAGCGCGTCAAAACTTCGCCTATGGGACGCAACATGATTACTTTGACC

ACCATAATATTATTGGATGGACACGCGAAGGGAATACCACACACCCCAAT

TCAGGATTAGCAACAATTATGTCGGACGGTCCAGGGGGCGAAAAATGGAT

GTATGTCGGACAAAACAAAGCAGGCCAAGTGTGGCATGACATAACAGGCA

ATAAACCGGGGACAGTGACGATTAACGCAGATGGCTGGGCAAATTTTTCA

GTCAACGGAGGTTCGGTCTCCATTTGGGTGAAAAGA

>Synthetic PrsA insert

ATGAAGAAGATTGCAATTGCGGCGATTACAGCGACAAGCGTGCTGGCTCT

CAGCGCATGCAGCGGGGGAGATTCTGAGGTTGTTGCGGAAACAAAAGCTG

GAAATATTACAAAAGAAGACCTTTATCAAACATTAAAAGACAATGCCGGA

GCGGACGCACTGAACATGCTTGTTCAGCAAAAAGTACTCGATGATAAATA

CGATGTCTCCGACAAAGAAATCGACAAAAAGCTGAACGAGTACAAAAAAT

CAATGGGTGACCAGCTCAACCAGCTCATTGACCAAAAAGGCGAAGACTTC

GTCAAAGAACAGATCAAATACGAACTTCTGATGCAAAAAGCCGCAAAGGA

TAACATAAAAGTAACCGATGATGACGTAAAAGAATATTATGACGGCCTGA

AAGGCAAAATCCACTTAAGCCACATTCTTGTGAAAGAAAAGAAAACGGCT

GAAGAAGTTGAGAAAAAGCTGAAAAAAGGCGAAAAATTCGAAGACCTTGC

AAAAGAGTATTCAACTGACGGTACAGCCGAAAAAGGCGGCGACCTCGGCT

GGGTCGGCAAAGACGATAACATGGACAAGGATTTCGTCAAAGCGGCATTT

GCTTTGAAAACCGGCGAAATCAGCGGACCTGTGAAATCCCAATTCGGCTA

TCACATCATTAAAAAAGACGAAGAACGCGGCAAATATGAAGACATGAAAA

AAGAGCTTAAAAAAGAAGTCCAAGAACAAAAGCAAAATGATCAAACTGAA

CTGCAATCCGTCATTGACAAACTTGTCAAAGATGCTGATTTAAAAGTAAA

AGACAAAGAGTTGAAAAAACAAGTCGACCAGCGTCAAGCTCAGACAAGCA

GCAGCAGC
